# Publication, Funding, and Experimental Data in Support of Human Reference Atlas Construction and Usage

**DOI:** 10.1101/2023.10.21.563417

**Authors:** Yongxin Kong, Katy Börner

## Abstract

Experts from 18 consortia are collaborating on the Human Reference Atlas (HRA) which aims to map the 37 trillion cells in the healthy human body. Information relevant for HRA construction and usage is held by experts (clinicians, pathologists, anatomists, single-cell experts), published in scholarly papers, and captured in experimental data. However, these data sources use different metadata schemes and cannot be cross-searched efficiently. This paper documents the compilation of a dataset, called HRAlit, that links the 136 HRA v1.4 digital objects (31 organs with 2,689 anatomical structures, 590 cell types, 1,770 biomarkers) to 583,117 experts; 7,103,180 publications; 896,680 funded projects, and 1,816 experimental datasets. The resulting HRAlit represents 23 tables with 21,704,001 records including 7 junction tables with 13,042,188 relationships. We demonstrate how HRAlit can be mined to identify leading experts, major papers, funding trends, or alignment with existing ontologies in support of systematic HRA construction and usage. Data and code are at https://github.com/cns-iu/hra-literature.

## Introduction

Constructing an atlas of the healthy human body is a massive undertaking due to the multiscale, biological complexity of human physiology. Since March 2020, international experts funded by the National Institutes of Health and/or supported by the Human Cell Atlas have been collaborating on the construction of a Human Reference Atlas (HRA)^1^. The 5th release of the HRA was published in June 2023 and comprises 31 organs with 2,689 unique anatomical structures, 590 unique cell types, 1,770 unique biomarkers linked to 31 Anatomical Structures, Cell Types, plus Biomarkers (ASCT+B) tables, 21 two-dimensional functional tissue units (FTU), and 65 three-dimensional, anatomically correct reference organs^2^. A total of 101 experts created and 99 experts reviewed (158 unique experts) the HRA digital objects across all releases and compiled 420 papers with DOIs that provide scholarly evidence for the anatomical structures, cell types, and biomarkers in the 31 ASCT+B tables.

As the HRA grows in the number of organs and data types it captures, it becomes important to use data-driven decision making to ensure systematic and efficient collaboration of scholars from different areas of research and development; federation of experimental data from different labs and data portals across scales (whole body to subcellular); and strategic foresight when setting data acquisition and tool development priorities.

In parallel to atlas construction, many high-quality experimental datasets are becoming available via data portals by HuBMAP^3^, SenNet^4^, KPMP^5,6^, GUDMAP^7^, GTEx^8^, or CZ CELLxGENE^9^. However, the portals use different metadata schemas and few use DOIs—cross-searching is difficult or impossible.

Moreover, HRA relevant data is published in scholarly papers. Each month, 88,777 papers are published in PubMed making it difficult to keep track of expertise, methods, data, code. *Scientific Data* papers typically focus on ontologies^10–12^ or experimental data^13–15^ while science of science studies commonly focus on authors, their publications, and possibly the funding that supports the research^16^. The construction of a HRA benefits from interlinking expertise, publication, funding, and experimental datasets.

This paper details the construction of the HRAlit database that links the 295 digital objects of the HRA (versions 1.0 to 1.4) to publication, funding, and experimental data in support of HRA construction and usage. Specifically, HRAlit includes 7,103,180 PubMed publications retrieved by a query for all 31 organs plus papers published in HRA, CZ CELLxGENE, GTEx, and CellMarker that are linked to 583,117 experts from 26,235 (cleaned) institutions and 896,680 funded projects by 6,427 (cleaned) funders. HRAlit also links the HRA to 1,816 experimental datasets and their 4,639 donors. The anatomical structures, cell types, and biomarkers in the 5th HRA release link to 5,049 ontology terms and IDs. The resulting database has 23 tables with 21,704,001 records including 7 junction tables with 13,042,188 relationships, including relationships between publications and organs, publications and datasets, publications and authors, authors and institutions, funding and funders (for detail, see **Table S1**).

Next, we demonstrate how HRAlit supports HRA construction and usage. For example, HRAlit (1) records what scholarly publication and experimental data evidence exists for which digital objects in the HRA; (2) makes it possible to identify experts that might like to serve as HRA authors and reviewers; (3) helps prioritize adding organs to the HRA for which sufficient experimental data and funding resources exist; (4) links HRA to existing ontologies so those can be expanded or used for ontological reasoning; (5) provides canonical data when exploring or annotating experimental data; and (6) supports analyses of HRA expertise and data diversity in terms of sex, ethnicity, geospatial coverage, etc..

## Results

### Interlinked Data

The HRAlit database covers seven data types (see **Table 1**) and their relationships in 23 tables (for detail of database descriptions and counts, see **Table S1**; for database schema, see **Fig. 1** and entity relationship diagram at https://github.com/cns-iu/hra-literature). It was constructed using these seven steps: (1) HRA data across five releases, including digital objects and experts, were downloaded, and the data from 5th release HRA, which includes publication references and ontology terms, was also acquired. (2) The names of all 31 organs covered in the 5th release HRA were used in a PubMed literature search, see Methods. (3) Dataset metadata and donor metadata (e.g., sex, age) for healthy human experimental data from HuBMAP, CZ CELLxGENE (CxG), and GTEx was downloaded. (4) Resulting publications were flagged as associated with HRA or experimental datasets, or listed in CellMarker for human cell makers and saved in the “*hralit_publication*” table together with their PMID, DOI, publication year, article title, and journal title. Citation and *h*-index metrics for publications were calculated from Web of Science. (5) All HRA experts and authors associated with selected publications, with ORCIDs listed in PubMed, were identified and saved in the “*hralit_author*” table that records ORCID, author first name, last name, earliest publication year, career age (see Methods), and total number of funded projects. Citation for publications and *h*-index metrics for authors were computed from Web of Science citation data. (6) Institution metadata was cleaned using OpenAlex. The cleaned institutions associated with author ORCIDs in the “*hralit_author*” table were stored in the “*hralit_institution*” table. (7) Funded projects listed in PubMed were stored in the “*hralit_funding*” table and the relationships among PMIDs, funding IDs, and funders sourced from PubMed were extracted and linked to the “*hralit_pub_funding_funder*” table. Funding organizations were cleaned using funder metadata and the relationships among PMID, funding IDs, and funders, as provided by OpenAlex. Funders associated with PMIDs and funded projects in the “*hralit_pub_funding_funder*” table were stored in the “*hralit_funder_cleaned*” table.

**Table 1.**
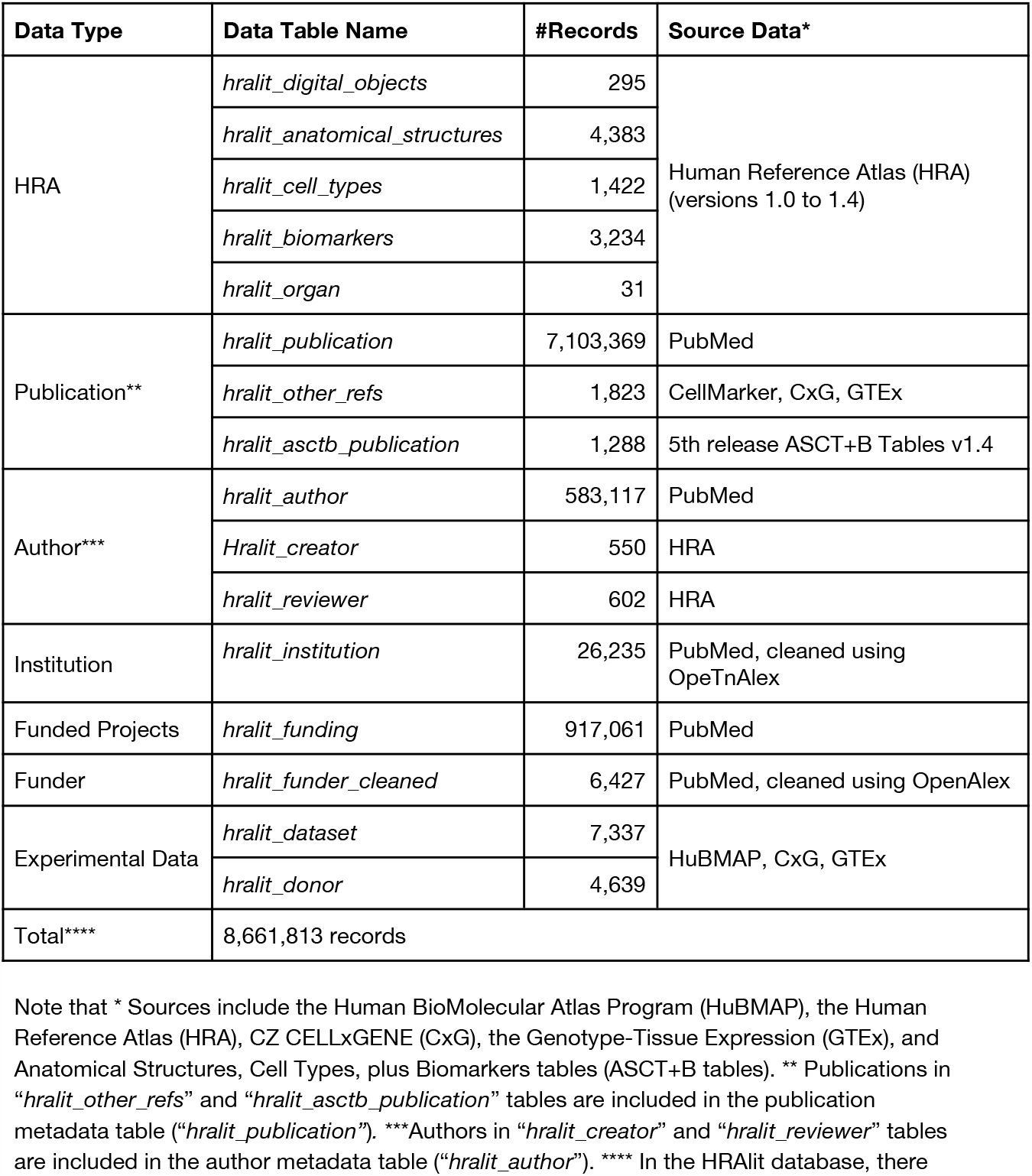
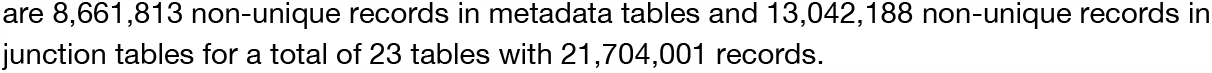
HRAlit data types, names, number of records, and source data.

| Data Type | Data Table Name | #Records | Source Data* |
| --- | --- | --- | --- |
| HRA | *hralit_digital_objects* | 295 | Human Reference Atlas (HRA)<br>(versions 1.0 to 1.4) |
|  | *hralit_anatomical_structures* | 4,383 |  |
|  | *hralit_cell_types* | 1,422 |  |
|  | *hralit_biomarkers* | 3,234 |  |
|  | *hralit_organ* | 31 |  |
| Publication** | *hralit_publication* | 7,103,369 | PubMed |
|  | *hralit_other_refs* | 1,823 | CellMarker, CxG, GTEx |
|  | *hralit_asctb_publication* | 1,288 | 5th release ASCT+B Tables v1.4 |
| Author*** | *hralit_author* | 583,117 | PubMed |
|  | *Hralit_creator* | 550 | HRA |
|  | *hralit_reviewer* | 602 | HRA |
| Institution | *hralit_institution* | 26,235 | PubMed, cleaned using OpeTnAlex |
| Funded Projects | *hralit_funding* | 917,061 | PubMed |
| Funder | *hralit_funder_cleaned* | 6,427 | PubMed, cleaned using OpenAlex |
| Experimental Data | *hralit_dataset* | 7,337 | HuBMAP, CxG, GTEx |
|  | *hralit_donor* | 4,639 |  |
| Total**** | 8,661,813 records |  |  |

Note that \* Sources include the Human BioMolecular Atlas Program (HuBMAP), the Human Reference Atlas (HRA), CZ CELLxGENE (CxG), the Genotype-Tissue Expression (GTEx), and Anatomical Structures, Cell Types, plus Biomarkers tables (ASCT+B tables). \*\* Publications in “*hralit\_other\_refs*” and “*hralit\_asctb\_publication*” tables are included in the publication metadata table (“*hralit\_publication*”). \*\*\*Authors in “*hralit\_creator*” and “*hralit\_reviewer*” tables are included in the author metadata table (“*hralit\_author*”). \*\*\*\* In the HRAlit database, there
are 8,661,813 non-unique records in metadata tables and 13,042,188 non-unique records in junction tables for a total of 23 tables with 21,704,001 records.

**Figure 1.**
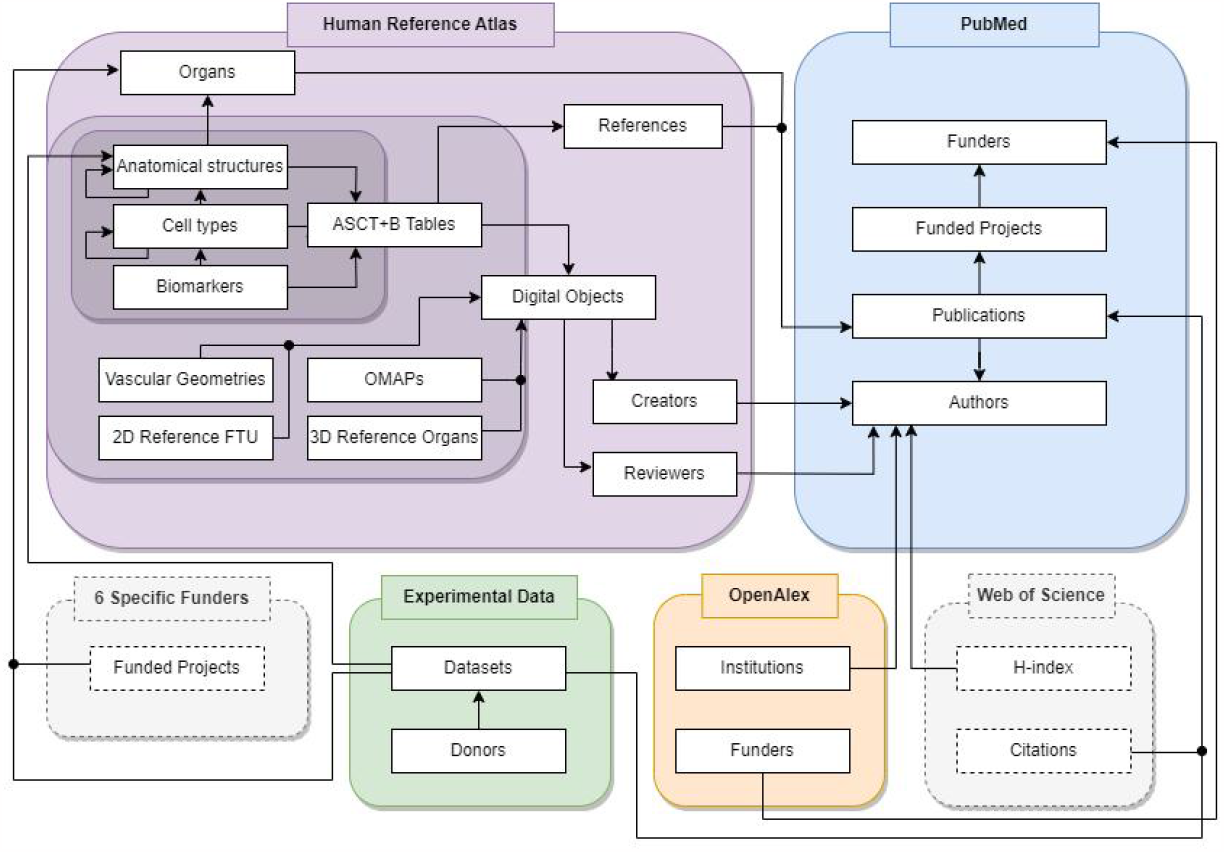
Overview of the HRAlit Database. Seven data types are interlinked by HuBMAP IDs of digital objects, PMIDs, DOIs, ORCIDs, funding IDs, dataset IDs, donor IDs, funder IDs, institution IDs. Note that WoS data is access restricted to CADRE affiliates; it cannot be shared publicly via the HRAlit. Additional funding data for six agencies Australian Research Data Commons (ARDC), the Canadian Institutes of Health Research (CIHR), European Commission (EC), Grants-in-Aid for Scientific Research (KAKEN), National Institutes of Health (NIH), and National Science Foundation (NSF) was downloaded for Fig 2c. but cannot be linked to HRAlit due to the differences in grant IDs used in PubMed and application IDs and funding project numbers in the funder datasets; it was not included in the HRAlit database.

All datasets and code used are detailed in the Methods section. Intermediate datasets and code used to retrieve, clean, and interlink the data can be found on GitHub at https://github.com/cns-iu/hra-literature. The HRAlit database is at https://github.com/cns-iu/hra-literature/blob/main/data/db/hralit.sql.

### Applications

Exemplarily, we present six applications that use the interlinked HRAlit database to (1) validate ASCT+B Tables using scholarly papers and experimental data evidence; (2)identify reviewers for HRA digital objects; (3) strategically decide what organs to focus on next for atlas construction; (4) connect the evolving HRA to diverse ontologies—in support of ontology updates but also ontology reasoning; (5) provide canonical markers for CellGuide, and (6) analyze the diversity and inclusiveness of HRA survey, HRA experts, general publication authors, and donor metadata.

### Providing Expert, Publication, and Experimental Data Evidence for the HRA

HRA authors and users need to understand what expertise, which publications, and what experimental data evidence exists for a specific organ. **Fig. 2** shows the number of experts, publications, funding (converted to U.S. dollars, see Methods), and experimental data for the 31 organs in the 5th release of the HRA (for details, see **Table S2**).

**Figure 2.**
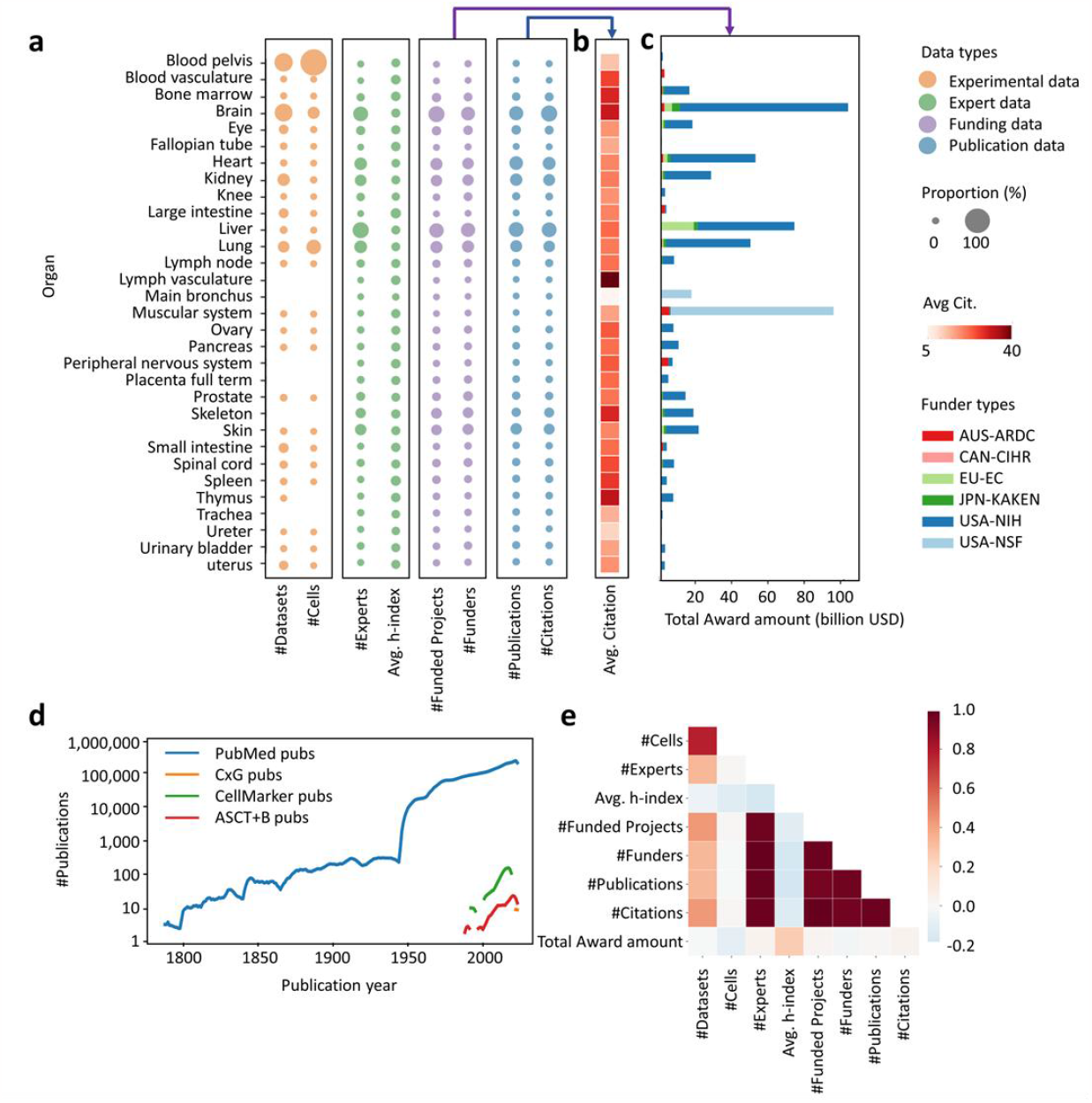
HRAlit Database and additional funding data. **a.** Experimental data, experts, funding, and publications for 31 organs in the HRA. Publication count refers to the number of publications (in hralit_publication). Dataset count refers to the number of experimental datasets per organ (in *hralit_dataset*). Funding count refers to the number of funded projects per organ that are cited by PubMed publications (in *hralit_funding*). Funder count refers to the number of funding agencies per organ that support the funding (in *hralit_funder_cleaned*). Expert count refers to the number of experts (authors and HRA experts) (*hralit_author*). Citation count and h-index count were retrieved from the Web of Science core collection. Total costs sum up funding by ARDC, CIHR, EC, KAKEN, NIH, NSF, converted to U.S. dollars using the latest annual exchange rate sourced from World Bank. Note that *hralit_institution* are not shown here. **b**. Average citation counts for 31 organs across 65% PubMed papers for which citation counts are available via WoS but not the 24,309,99 other papers. **c**. Total award amount from six funders in four countries and European Union. **d**. The trend of publication output related to the 31 organs. **e**. Correlation between different HRA evidence, e.g., the number of datasets and total cells but also the number of experts and publications are both positively correlated.

Note that average citation counts differ across organs—analogous to differences in citation counts that are typical in scholarly disciplines^16,17^. For example, research on main bronchus receives comparatively little funding and generates about 8 papers per year that have a comparatively low number of citations—about 3 citations on average, with the highest cited paper (entitled “Induced bronchus-associated lymphoid tissue serves as a general priming site for T cells and is maintained by dendritic cells”) having 183 citations. In contrast, brain research enjoys a total of 20 billion U.S. dollar in funding as acknowledged in PubMed, generates 4,825 papers each year, and there are 32 citations per paper on average with the highest cited paper entitled “Analysis of relative gene expression data using real-time quantitative PCR and the 2(-Delta Delta C(T)) Method” having 106,059 citations.

### Identify HRA Reviewers

The 5th release of the HRA comprises 136 digital objects (DO) including 31 ASCT+B Tables plus a crosswalk, 65 3D reference objects plus two united files (one for male and another for the female body; each comprising all organs) and a crosswalk, 21 2D reference objects plus a crosswalk, 13 Organ Mapping Antibody Panels (OMAP) objects, and one Vascular Common Coordinate Framework (VCCF) object. Across all five HRA releases, there are 295 digital objects. All of these DOs need to be reviewed by domain experts using a process analogous to reviewing scholarly papers before publication. The HRAlit database was used to identify experts with highly cited papers, funding, and expertise for a certain organ. Title and MeSH terms assigned to 31 organs associated with papers published by an expert author were used to predict the expertise subjects (e.g., anatomy and pathology expertise is needed to review anatomical structures while single cell and chemistry expertise is needed to review cell types and/or biomarkers in the ASCT+B Tables). A total of 583,117 unique experts with ORCIDs were retrieved, including both HRA experts and authors associated with all selected publications. Many of them have expertise in multiple organs, and those with high citation counts and many funded projects were invited to serve as reviewers. Exactly 34 experts agreed, served as reviewers, and are/will be acknowledged in future releases of the HRA.

### Prioritize Atlas Construction

Single-cell data experiments are expensive—acquiring gene or protein expression data costs about 30 cents per cell. A typical tissue sample has about 10,000 cells and costs about 300,000 U.S. dollars on average. Many datasets and different assay types are required to construct an atlas. The HRAlit database can be used to identify the number of existing datasets per organ and the likely future number of datasets based on available funding. Plus, the number of available experts matters as human expertise is key for atlas construction as are scholarly publications that provide detailed documentation for high quality datasets. **Table 2** lists the types of DOs and HRAlit data types per organ and it is actively used to prioritize HRA construction.

**Table 2.**
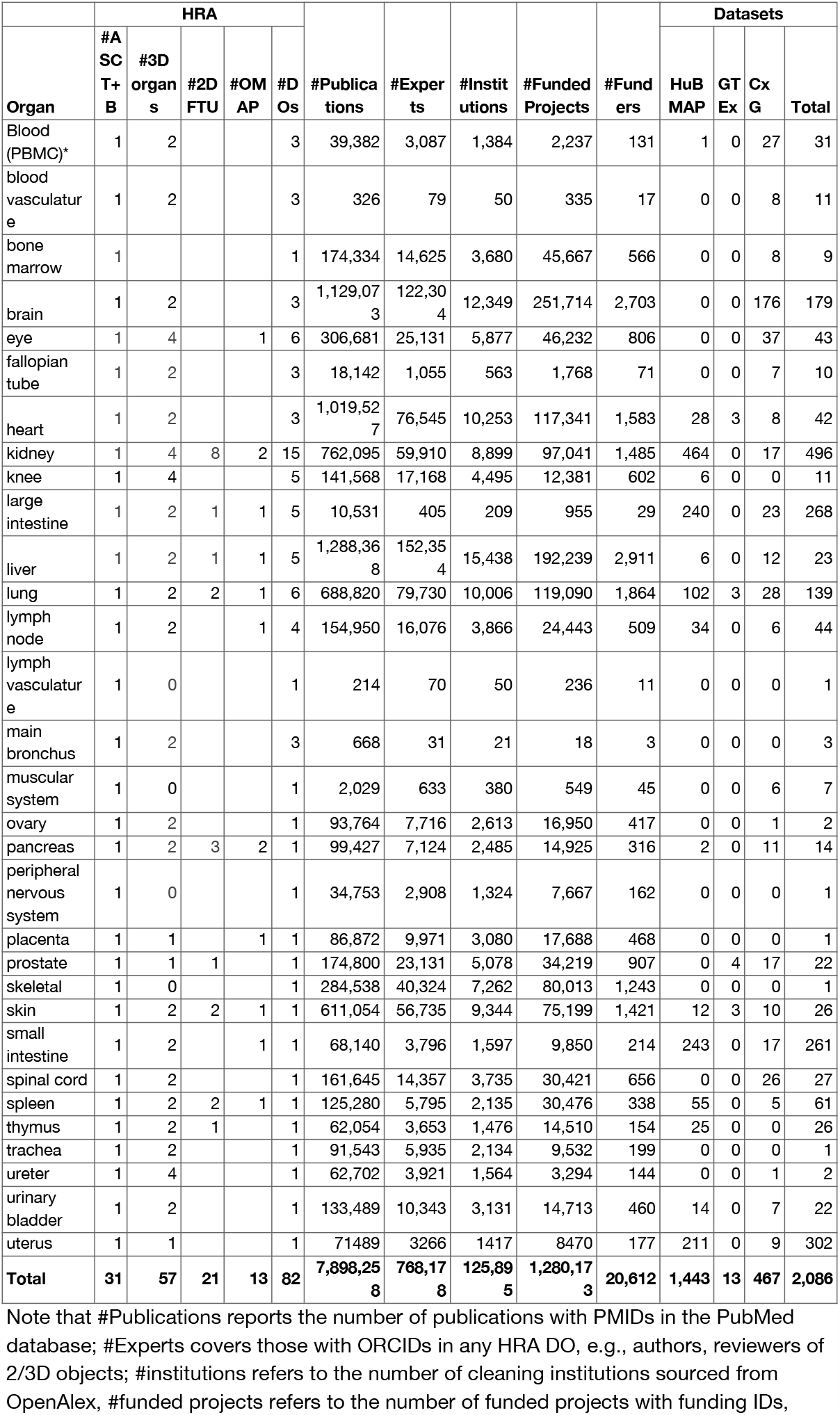
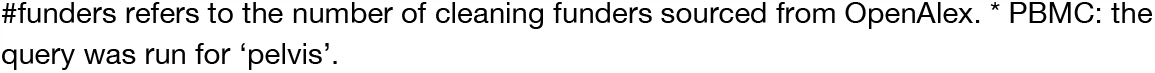
Publication evidence, experts, institutions, funded projects, funders, and experimental data available for 31 organs. Rows represent the organs from the 5th HRA release (bronchus is part of the lung). For each organ, we provide the number of HRA DO types and the number of publications, experts, institutions, funded projects and funders, together with the number of healthy human datasets available via the HuBMAP, GTEx, and CELLxGENE portals on Aug 1, 2023.

| Organ | HRA |  |  |  |  | #Publications | #Experts | #Institutions | #Funded Projects | #Funders | Datasets |  |  |  |
| --- | --- | --- | --- | --- | --- | --- | --- | --- | --- | --- | --- | --- | --- | --- |
|  | #A SC T+B | #3D organs | #2D FTU | #OM AP | #D Os |  |  |  |  |  | HuB MAP | GT Ex | Cx G | Total |
| Blood (PBMC)* | 1 | 2 |  |  | 3 | 39,382 | 3,087 | 1,384 | 2,237 | 131 | 1 | 0 | 27 | 31 |
| blood vasculature | 1 | 2 |  |  | 3 | 326 | 79 | 50 | 335 | 17 | 0 | 0 | 8 | 11 |
| bone marrow | 1 |  |  |  | 1 | 174,334 | 14,625 | 3,680 | 45,667 | 566 | 0 | 0 | 8 | 9 |
| brain | 1 | 2 |  |  | 3 | 1,129,073 | 122,304 | 12,349 | 251,714 | 2,703 | 0 | 0 | 176 | 179 |
| eye | 1 | 4 |  | 1 | 6 | 306,681 | 25,131 | 5,877 | 46,232 | 806 | 0 | 0 | 37 | 43 |
| fallopian tube | 1 | 2 |  |  | 3 | 18,142 | 1,055 | 563 | 1,768 | 71 | 0 | 0 | 7 | 10 |
| heart | 1 | 2 |  |  | 3 | 1,019,527 | 76,545 | 10,253 | 117,341 | 1,583 | 28 | 3 | 8 | 42 |
| kidney | 1 | 4 | 8 | 2 | 15 | 762,095 | 59,910 | 8,899 | 97,041 | 1,485 | 464 | 0 | 17 | 496 |
| knee | 1 | 4 |  |  | 5 | 141,568 | 17,168 | 4,495 | 12,381 | 602 | 6 | 0 | 0 | 11 |
| large intestine | 1 | 2 | 1 | 1 | 5 | 10,531 | 405 | 209 | 955 | 29 | 240 | 0 | 23 | 268 |
| liver | 1 | 2 | 1 | 1 | 5 | 1,288,368 | 152,354 | 15,438 | 192,239 | 2,911 | 6 | 0 | 12 | 23 |
| lung | 1 | 2 | 2 | 1 | 6 | 688,820 | 79,730 | 10,006 | 119,090 | 1,864 | 102 | 3 | 28 | 139 |
| lymph node | 1 | 2 |  | 1 | 4 | 154,950 | 16,076 | 3,866 | 24,443 | 509 | 34 | 0 | 6 | 44 |
| lymph vasculature | 1 | 0 |  |  | 1 | 214 | 70 | 50 | 236 | 11 | 0 | 0 | 0 | 1 |
| main bronchus | 1 | 2 |  |  | 3 | 668 | 31 | 21 | 18 | 3 | 0 | 0 | 0 | 3 |
| muscular system | 1 | 0 |  |  | 1 | 2,029 | 633 | 380 | 549 | 45 | 0 | 0 | 6 | 7 |
| ovary | 1 | 2 |  |  | 1 | 93,764 | 7,716 | 2,613 | 16,950 | 417 | 0 | 0 | 1 | 2 |
| pancreas | 1 | 2 | 3 | 2 | 1 | 99,427 | 7,124 | 2,485 | 14,925 | 316 | 2 | 0 | 11 | 14 |
| peripheral nervous system | 1 | 0 |  |  | 1 | 34,753 | 2,908 | 1,324 | 7,667 | 162 | 0 | 0 | 0 | 1 |
| placenta | 1 | 1 |  | 1 | 1 | 86,872 | 9,971 | 3,080 | 17,688 | 468 | 0 | 0 | 0 | 1 |
| prostate | 1 | 1 | 1 |  | 1 | 174,800 | 23,131 | 5,078 | 34,219 | 907 | 0 | 4 | 17 | 22 |
| skeletal | 1 | 0 |  |  | 1 | 284,538 | 40,324 | 7,262 | 80,013 | 1,243 | 0 | 0 | 0 | 1 |
| skin | 1 | 2 | 2 | 1 | 1 | 611,054 | 56,735 | 9,344 | 75,199 | 1,421 | 12 | 3 | 10 | 26 |
| small intestine | 1 | 2 |  | 1 | 1 | 68,140 | 3,796 | 1,597 | 9,850 | 214 | 243 | 0 | 17 | 261 |
| spinal cord | 1 | 2 |  |  | 1 | 161,645 | 14,357 | 3,735 | 30,421 | 656 | 0 | 0 | 26 | 27 |
| spleen | 1 | 2 | 2 | 1 | 1 | 125,280 | 5,795 | 2,135 | 30,476 | 338 | 55 | 0 | 5 | 61 |
| thymus | 1 | 2 | 1 |  | 1 | 62,054 | 3,653 | 1,476 | 14,510 | 154 | 25 | 0 | 0 | 26 |
| trachea | 1 | 2 |  |  | 1 | 91,543 | 5,935 | 2,134 | 9,532 | 199 | 0 | 0 | 0 | 1 |
| ureter | 1 | 4 |  |  | 1 | 62,702 | 3,921 | 1,564 | 3,294 | 144 | 0 | 0 | 1 | 2 |
| urinary bladder | 1 | 2 |  |  | 1 | 133,489 | 10,343 | 3,131 | 14,713 | 460 | 14 | 0 | 7 | 22 |
| uterus | 1 | 1 |  |  | 1 | 71489 | 3266 | 1417 | 8470 | 177 | 211 | 0 | 9 | 302 |
| **Total** | **31** | **57** | **21** | **13** | **82** | **7,898,258** | **768,178** | **125,895** | **1,280,173** | **20,612** | **1,443** | **13** | **467** | **2,086** |

Note that #Publications reports the number of publications with PMIDs in the PubMed database; #Experts covers those with ORCIDs in any HRA DO, e.g., authors, reviewers of 2/3D objects; #institutions refers to the number of cleaning institutions sourced from OpenAlex, #funded projects refers to the number of funded projects with funding IDs,

We also list the availability of OMAP-aligned spatial proteomics data^18^ that can be easily mapped to the HRA. Also listed is the availability of single cell bulk data annotation tools such as Azimuth^19^, Cell Typist^20^ recently renamed to CellHint^21^, or popV used in Tabula Sapiens^22^.

### Linking the HRA to Existing Ontologies

The 5th release of the HRA covers (1) 2,691 HRA anatomical structures that link to 1,443 terms in Uberon and 1,246 in FMA; (2) the 590 cell types link to 127 terms in PCL, 461 in CL and 621 new ASCTB-TEMP that do not yet exist in ontologies, and 1 term in LMHA; (3) 1,770 biomarkers, such as HGNC, GNC, LOC, UNIPROT, LM, and PR, which associated with 1,608 genes, 460 proteins, and 1 lipids. CellMarker 2.0^23^ is a manually curated database of experimentally supported gene and protein biomarkers for cell types in humans and mice. Here, we compare 31 organs covered in the HRA to those covered in the human part of CellMarker 2.0 and CZ CELLxGENE (see **Table 3**).

**Table 3.**
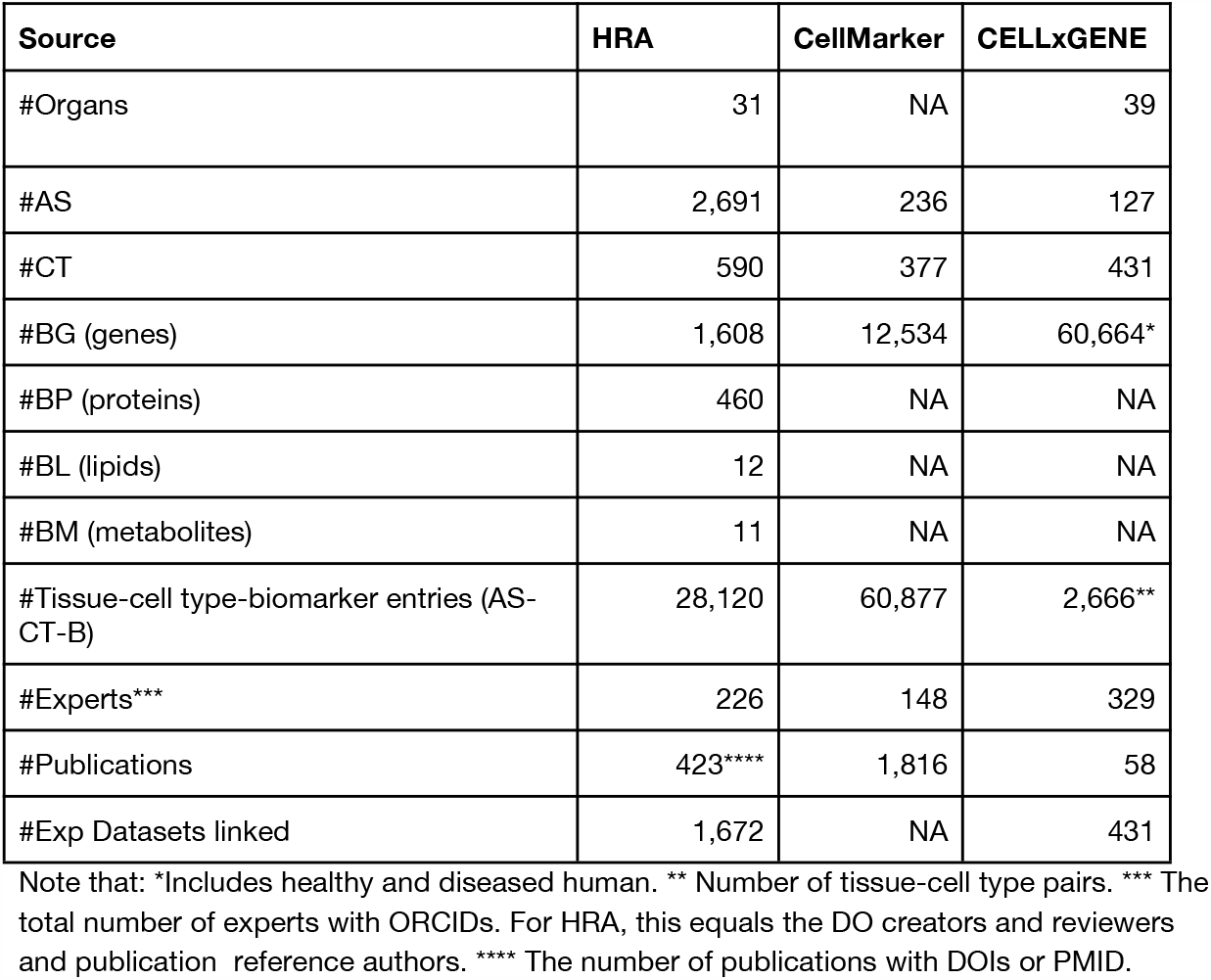
Comparing HRA, CellMarker, CELLxGENE for 31 healthy human adult organs. Note that HRA reports gene, protein, lipid, and metabolite biomarkers; CellMarker focuses on tissue-cell type-biomarker triples; CELLxGENE focuses on gene biomarkers.

| Source | HRA | CellMarker | CELLxGENE |
| --- | --- | --- | --- |
| #Organs | 31 | NA | 39 |
| #AS | 2,691 | 236 | 127 |
| #CT | 590 | 377 | 431 |
| #BG (genes) | 1,608 | 12,534 | 60,664* |
| #BP (proteins) | 460 | NA | NA |
| #BL (lipids) | 12 | NA | NA |
| #BM (metabolites) | 11 | NA | NA |
| #Tissue-cell type-biomarker entries (AS-CT-B) | 28,120 | 60,877 | 2,666** |
| #Experts*** | 226 | 148 | 329 |
| #Publications | 423**** | 1,816 | 58 |
| #Exp Datasets linked | 1,672 | NA | 431 |

Note that: \*Includes healthy and diseased human. \*\* Number of tissue-cell type pairs. \*\*\* The total number of experts with ORCIDs. For HRA, this equals the DO creators and reviewers and publication reference authors. \*\*\*\* The number of publications with DOIs or PMID.

### Compare Canonical HRA Marker Genes with Computational CZI Genes in CellGuide

CZ CELLxGENE^9^ recently added the 5th release of the HRA data to the CellGuide tool, see **Fig. 3**. CellGuide helps researchers understand what cells in the Cell Ontology (https://www.ebi.ac.uk/ols4/ontologies/cl) are in the experimental dataset corpus (blue coloured nodes) and which are not (grey nodes). Subtrees can be expanded to see descendant nodes.

**Figure 3.**
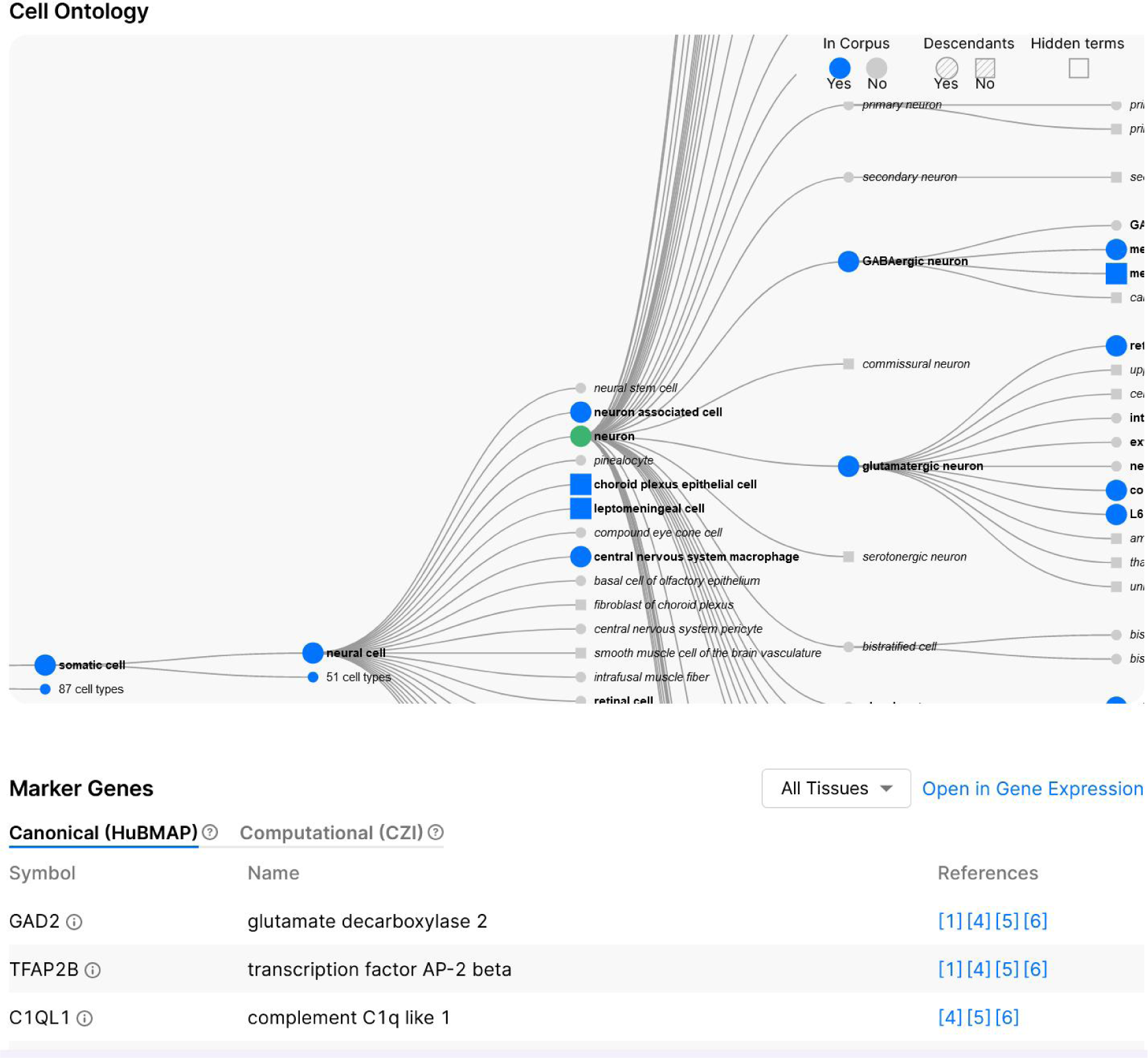
CZ CellGuide visualization for Neuron (CL:0000540) showing the CL ontology typology with the ‘neuron’ cell type highlighted in green together with its parent (‘neural cell’, which *is_a* ‘somatic cell’, which *is_a* ‘animal cell’) and children nodes (e.g., ‘GABAergic neuron’ and ‘gutamagergic neuron’). The interactive visualization is at https://cellxgene.cziscience.com/cellguide/CL_0000540.

On Aug 3, 2023, there are 2,706 cell types in CL, interlinked by *is_a* child to parent relations; the root note of this cell typology is ‘cell’. Plus, there are 791 cell types (613 for homo sapiens) in the CxG experimental data and 296 of these cell types are also in the 5th release ASCT+B Tables. Note that CellGuide only features cell types from the ASCT+B Tables that are connected to the CL typology—omitting the 621 ASCTB-TEMP and 1 term in the LungMAP Human Anatomy (LMHA) that do not yet exist in CL. A total of 588 cell types (127 in PCL and 461 in CL) are connected via *is_a* relationships but only cell types that are in the CxG experimental data are shown in CellGuide.

In CellGuide, a cell type is associated with the most discriminating gene biomarkers across all anatomical structures in the human body. Cell markers can be subset by organ using the ‘All Tissues’ menu (see **Fig. 3**) or by organism: For example, for ‘T cell’ (https://cellxgene.cziscience.com/cellguide/CL_0000084), if users navigate to ‘Marker Genes’ and then selects the ‘Computational (CZI)’ tab, they can select ‘Homo sapiens’, ‘Macaca mulatta’ or ‘Mus musculus’.

CellGuide accesses organ (in CZ CELLxGENE called ‘Tissue’) specific gene biomarkers via the ASCT+B API (https://mmpyikxkcp.us-east-2.awsapprunner.com/#/). References were parsed out of the ASCT+B Table CSV files. The 6th release of the HRA API, out in December 2023, will make it easier to access this data via API queries.

Using CellGuide, experts can compare cell types and marker genes in the CZ CELLxGENE experimental data—called Computational (CZI)—to marker genes in the human authored ASCT+B Tables—Canonical (HuBMAP)—and HRA publication references.

CellGuide is valuable for improving the coverage and quality of ASCT+B Tables in future HRA iterations. ASCT+B Table authors are provided with a listing of cell types and gene biomarkers seen in healthy adult data and are asked to consider inclusion of these cell types and biomarkers in future table iterations.

### Enhancing Diverse Perspectives

The HRA is meant to present healthy humans in an inclusive manner. Ideally, the demographics of experts engaged and human tissue data used matched the demographic of the world population. **Fig. 4** and **Table S3** report gender and (career) age via population pyramids, ethnic composition of experts and donor data, and geolocation for experts engaged in and datasets used for HRA construction.

**Figure 4.**
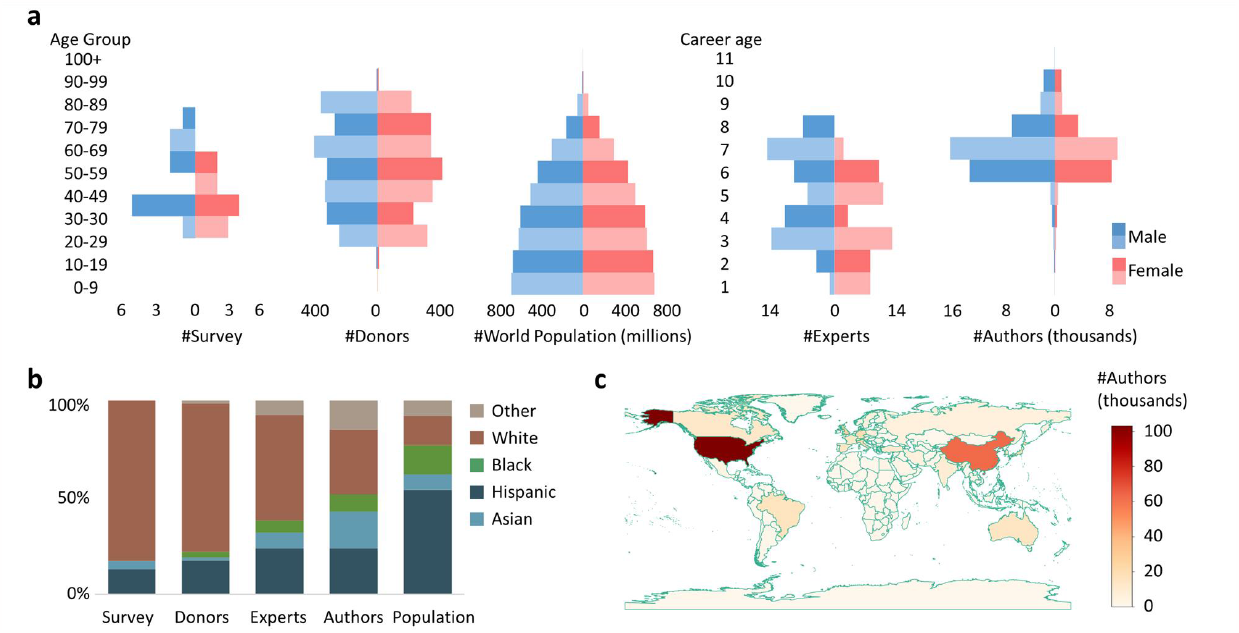
Comparison of HRA, publication, and experimental to world population data. **a.** Population pyramids by age group of survey respondents and tissue data donors in comparison to world population. Population pyramids by career age for HRA experts and publication authors. **b**. Ethnic composition of survey respondents, HRA tissue donors, HRA experts, paper authors, world population in percentages. **c**. Choropleth map showing the number of paper authors overlaid on a world map.

The data comes from six different sources:

1. A survey run annually to capture the diversity of experts engaged in the monthly Human Reference Atlas working group meetings; April 2023 survey responses by 23 respondents are shown here.
2. Experts listed as authors and reviewers of the digital objects in the 5th HRA release at https://hubmapconsortium.github.io/ccf-releases/v1.4/docs.
3. Authors of publications listed in the ASCT+B Tables in the 5th HRA release.
4. Authors of the publications and funded projects retrieved from PubMed, CellMarker, and experimental datasets for the 31 organs in the 5th release HRA.
5. Donor metadata for experimental datasets served via the HuBMAP, GTEx, and CZ CELLxGENE data portals.
6. World population data for 2023 from PopulationPyramid portal downloaded from https://www.populationpyramid.net/api/pp/900/2023/?csv=true.

The gender of authors was predicted using the gender_guesser package, which tags gender based on first names. The race of authors was determined using the ethnicolr package, based on both first and last names. Career age was estimated based on the publication year of an author’s first paper, as detailed in the Methods section.

## Discussion

The paper presented the compilation of a unique database that considerably extends the scope and value of the HRA. Specifically, we linked HRA data to publications, experts and their institutions, funded projects, and funders, experimental data and ontologies. The value of the HRAlit database was showcased in six applications that aim to improve the quality, coverage, utility, and inclusiveness of the HRA. All code used to compile the data was made available on GitHub in October 2023, and will be re-run for each HRA release every six months.

There are several known data limitations. First, some organs have multiple names but only their official ontology term was used to retrieve PubMed publications (e.g., “large intestine” but not “colon” was used as a query term) and no synonyms were used. Second, the CADRE provided WoS database is no longer being updated; the current copy covers years 1898 to 2023. Third, we plan to add data from EBI’s Single Cell Expression Atlas (https://www.ebi.ac.uk/gxa/sc)^24,25^. Cellpedia (https://cellpedia.org/en/cell-therapy) and CellKb (https://www.cellkb.com) do not currently offer APIs that support data download but would add value.

Going forward, we will link the HRAlit database to the Scalable Precision Medicine Open Knowledge Engine (SPOKE) biomedical knowledge graph^26,27^ that interlinks 27 million concepts (21 types, including genes, proteins, diseases) via 53 million semantically meaningful relationships (55 types, including TREATS_CtD and CONTRAINDICATES_CcD connecting compounds and diseases) downloaded from 41 databases that use 11 ontologies (including Uberon, CL). This expansion of HRAlit will support novel queries. For example, SPOKE has Gene-encodes-Protein assertions that can be mined to compare gene biomarkers from single cell bulk assays with protein biomarkers from spatial assays. SPOKE also links molecular to cellular entities to pharmacology and clinical practice making it possible to run knowledge graph queries in support of drug discovery and precision health.

## Methods

### Data

An overview of the HRAlit database is given in **Fig. 1** and in the entity relationship diagram at https://github.com/cns-iu/hra-literature).

### Human Reference Atlas

**HRA digital objects** for the HRA 5th release (v1.4) are listed at https://hubmapconsortium.github.io/ccf-releases/v1.4/docs and include 31 ASCT+B Tables plus 1 crosswalk table, 21 Functional Tissue Units (FTUs), 13 Organ Mapping Antibody Panels (OMAPs), 1 Vascular Geometries table, as well as 65 3D Reference Organs plus 3 digital objects for two united and 1 crosswalk from the 3D organs to the ASCT+B tables for a total of 136 digital objects. All 295 HRA digital objects across all HRA releases (starting with v1.0 in March 2021 to v1.4 in June 2023) can be accessed via the CCF-HRA releases^28^, and extracted as a HRA metadata table. The 5th release ASCT+B tables are available at https://ccf-ontology.hubmapconsortium.org/v2.2.1/ccf-asctb-all.json. The linkages among anatomical structures, cell types, and biomarkers are extracted from the 5th release ASCT+B tables. In total, there are 4,835 *part_of* edges between 4,384 unique anatomical structures, 7,879 *located_in* edges between 691 unique cell types and these anatomical structures, 620 *is_a* linkages between cell types, and 6,812 *characterizes* edges between 2,927 unique biomarkers and the unique cell types.

### HRA Experts

Information about the creators and reviewers of HRA across all versions is selected based on the HRA metadata table. Resulting table “*hralit_creator*” and “*hralit_reviewer*” includes ORCIDs, first names, and last names, associated digital objects. In total, there are 101 unique creators and 99 unique reviewers for a total of 158 unique experts.

### HRA References

Each publication in the 5th release ASCT+B tables is characterized by an ID, DOI, accompanying notes, categorization type, and an associated organ. Within the HRA reference framework, publication references are classified as “*general_publication*” that supports organ-specific ASCT-B tables and are listed in the table header and “*reference*” papers that support combinations of table row-specific Anatomical Structures (AS), Cell Types (CT), and Biomarkers (B). In total, the 5th release ASCT+B tables has 83 general references and 1,070 specific references for a total of 1,057 unique publication references. 420 of these publications have DOIs and 391 are in PubMed.

### HRA Ontologies

Anatomical structures, cell types, and biomarkers in the HRA are linked to their counterparts in existing ontologies; for 621 terms there do not exist entries in ontologies and HRA TEMP IDs are assigned to facilitate ID-based mapping while these new terms are added to existing ontologies. Ontology IDs were extracted via 5th release ASCT+B tables (https://ccf-ontology.hubmapconsortium.org/v2.2.1/ccf-asctb-all.json). In total, 1,246 anatomical structures are linked to the Foundational Model of Anatomy (FMA) and 1,443 to the Uber-anatomy ontology (Uberon), 127 cell types are linked to entries in the Cell Ontology (CL) and 127 Provisional Cell Ontology (PCL), 1,669 biomarkers are linked to the Human Genome Nomenclature Committee (HGNC), and 101 are linked to another biomarker ontology (e.g., UniProt). Note that there are issue requests for the 621 ASCTB-TEMP that do not yet exist in CL; these will be added over the coming 1-2 years to properly represent healthy human cells in CL.

**HRA Diversity survey data** is available at https://github.com/cns-iu/hra-literature.

### CellMarker

CellMarker 2.0^23^ is a manually curated dataset of biomarkers for distinguishing different cell types in different anatomical structures in human and mouse derived from and linked to over 100,000 publication references. For human, CellMarker covers 13,605 biomarkers, 467 cell types, and 158 anatomical structures linked to ontologies such as Uberon, CL, NCBI Gene, and UniProt. Data was downloaded manually as a file from the CellMarker portal at http://xteam.xbio.top/CellMarker, see counts in **Table 3**. Publications associated with CellMarker are tagged as “cellmarker” in HRAlit and are stored in the “*hralit_other_refs*” table, including 1,622 publications with DOIs.

### Publications

PubMed (https://pubmed.ncbi.nlm.nih.gov) provides free access to publications encompassing various health and life sciences disciplines. Daily updates of PubMed can be downloaded via API (https://www.ncbi.nlm.nih.gov/books/NBK25497/). We queried a local, daily updated database on September 12, 2023 using 31 organ names as query keywords. 7,103,180 unique publications covering years 1900 to 2023 (see **Table S2** for the number of papers per publication year) resulted and the HRAlit database covers their identifiers (PMIDs and DOIs), article titles, journal titles, publication year, authors, among others.

Web of Science (WoS) (https://www.webofscience.com/wos) provides citations for publications. WoS XML raw data was acquired by Indiana University Network Institution from Clarivate Analytics, and the Cyberinfrastructure for Network Science Center has access to this data under a Data Custodian user agreement the Collaborative Archive & Data Research Environment (CADRE)^30^. Based on the linkages between WoS IDs and PMIDs provided by WoS, citations are counted as the total number of papers that cited a publication.

### Experts

Author details were extracted from PubMed using unique identifiers for authors, known as ORCID IDs. There are 583,117 authors with ORCID identifiers that authored 483,998 of the 7,103,180 publications. The HRAlit database covers for each author: ORCID IDs, first name, last name, publication year of author’s first publication, career age, and the number of unique funded projects assigned to publications authored by an author, which is stored in the “*hralit_author*” table. Additionally, HRAlit database also includes linkages associated with authors, with author-institutions linkages stored in the “*hralit_author_institution*” table, author-publication linkages in the “*hralit_publication_author*” table, author-organ linkages in the “*hralit_author_expertise*” table. Based on the publication citations, *h*-index is also calculated through the linkage between PMIDs and ORCIDs.

### Institutions

Author institutions provided by PubMed were cleaned using data from the OpenAlex database^31^ on August 31, 2023. OpenAlex sources institution data from Crossref, PubMed, ROR, MAG, and various publisher websites, focusing on indexing institutions associated with authors. Accessible via the OpenAlex API (https://docs.openalex.org), each institution is uniquely identified using both an internal ID (e.g., “https://openalex.org/I2802101240“) and a canonical external ID, called ROR ID. OpenAlex provides the institution name, institution type, and country code for each registered entity. We obtained 102,494 institution entities from OpenAlex, of which 26,235 institutions linked to 463,617 unique ORCIDs covering 79.50% of 583,117 unique ORCIDs, see “*hralit_institution*” table in the HRAlit database.

## Experimental Data

Tissue dataset metadata was downloaded from three public data portals on Aug 7, 2023.

**HuBMAP Portal** (https://portal.hubmapconsortium.org): The HuBMAP SmartAPI was used to download 1,610 tissue samples from 193 donors with 1,445 associated datasets using https://entity.api.hubmapconsortium.org/datasets/prov-info. Note that there are many more datasets that are currently in preprocessing and quality control stages and will soon become available; with member-login, there are 411,200 tissue samples from 397 donors with 3,628 associated datasets.

**CZ CELLxGENE Portal** (https://cellxgene.cziscience.com): Using the cellxgene-census API at https://chanzuckerberg.github.io/cellxgene-census/index.html, 593 datasets from 5,328 donors, linked to 58 publications with DOIs, were downloaded.

**GTEx Portal** (https://gtexportal.org/home): Single cell data for 25 samples from 16 donors was downloaded manually from https://www.science.org/doi/suppl/10.1126/science.abl4290/suppl_file/science.abl4290_tables_s1_to_s20.zip^31^.

### Table Schema for HRAlit Database

The HRAlit database, with a total size of 1.56 GB, is available in SQL format at https://github.com/cns-iu/hra-literature. Additionally, each table is also provided in CSV format. For a detailed data dictionary see **Table S1**.

### Algorithms

#### Data extraction and preprocessing

HRA information on digital objects, creators, and reviewers is extracted from the HRA metadata. Ontology IDs and HRA publication references are extracted from ASCT+B Tables.

Donor and other metadata for experimental datasets from HuBMAP and CxG were downloaded via the HuBMAP and CxG APIs. GTEx single cell data was derived from the supplementary tables of Eraslan et al^31^. The final datasets and donors are integrated using a manually curated dataset mapping table.

PubMed publications, citations, authors, funding, and funders data were retrieved from a daily updated PostgreSQL database using PSQL queries for 31 organ names in titles and MeSH terms. Authors with ORCIDs were extracted and tagged with the same organ labels as their associated publications. Finally, funding and funder data were extracted from the publications.

Using the OpenAlex API, metadata related to institutions and funders was obtained. This also includes the relationships between institutions and authors, as well as between funders, funding, and publications, with a focus on PubMed publications. To clean the institution and funding agency data, institution IDs are linked to the selected authors based on matching author ORCIDs. Similarly, funder IDs are connected to selected publications and funding from PubMed based on congruent PMIDs and funding IDs.

For **Fig. 2** and **Table S2** but not for use in HRAlit, we obtained citation data from Web of Science (WoS) through queries on a PostgreSQL database provided by the Collaborative Archive & Data Research Environment (CADRE) project; funding data for six specific funders was extracted using data portal APIs.

#### Database construction

To create the HRAlit database, follow these steps: (1) Load data to create all 17 data tables. (2) Join data. Data from 17 tables is combined based on related columns to form 6 comprehensive tables. (3) Link data. Relationships between tables are established, ensuring data integrity and enforcing referential constraints. (4) Export database. HRAlit database is exported in both SQL and CSV formats.

#### Diversity - Predict Gender, Ethnicity

Gender is predicted using the “gender_guesser” Python module (https://pypi.org/project/gender-guesser), which classifies authors based on first names into categories including “male”, “female”, “andy”, “mostly_male”, “mostly_female”, “unknown”. For classifications other than “male” or “female” for 36 experts, gender was identified by examining photos found on Google using full names with affiliated institutions from ORCID, with exception of one expert named Li Yao, who lacked affiliation information. Ethnicity is determined using the “ethnicolr” Python module, which is trained on U.S. Census data and other large-scale name-ethnicity datasets. Based on both first names and last names, the module calculates probabilities for five ethnic categories: Asian, Black, Hispanic, White, and Other.

## Data Availability

All data and an entity relationship diagram are available at https://github.com/cns-iu/hra-literature.

## Code Availability

All code for data extraction, database construction, and applications is available at https://github.com/cns-iu/hra-literature. Code is licensed under the MIT License and data under the Creative Commons Attribution 4.0 International License.

## Acknowledgements

We thank Bruce W. Herr II for collecting metadata and linkages from OpenAlex; Jason Hilton, Brian Raymor, and Alexander Tarashansky for offering guidance on selecting data from CellGuide; Mike Gallant and Ben Serrette for providing access to the PubMed data and Web of Science data from Indiana University Network Science through the Collaborative Archive & Data Research Environment (CADRE) project; Nancy Ruschman and Michael Ginda for their expert comments on an earlier version of this paper.

This research has been funded by the China Scholar Council [YK] and the NIH Common Fund through the Office of Strategic Coordination/Office of the NIH Director under awards OT2OD033756 and OT2OD026671, by the Cellular Senescence Network (SenNet) Consortium through the Consortium Organization and Data Coordinating Center (CODCC) under award number U24CA268108, by the Kidney Precision Medicine Project grant U2CDK114886, by the NIDDK under awards U24DK135157 and U01DK133090 and by The Multiscale Human CIFAR project [KB]. The funders had no role in study design, data collection and analysis, decision to publish, or preparation of the manuscript.

## Author contributions

YK compiled all data, constructed the HRAlit database, ran the data analyses and visualizations, and co-wrote the paper.

KB led the design and implementation of the study and co-wrote the paper.

