## Supplementary Tables for "Publication, Funding, and Experimental Data in Support of Human Reference Atlas Construction and Usage"

Experts from 18 consortia are collaborating on the Human Reference Atlas (HRA) which aims to map the 37 trillion cells in the healthy human body. Information relevant for HRA construction and usage is held by experts (clinicians, pathologists, anatomists, single-cell experts), published in scholarly papers, and captured in experimental data. However, these data sources use different metadata schemes and cannot be cross-searched efficiently. This paper documents the compilation of a dataset, called HRAlit, that links the 136 HRA v1.4 digital objects (31 organs with 2,689 anatomical structures, 590 cell types, 1,770 biomarkers) to 583,117 experts; 7,103,180 publications; 896,680 funded projects, and 1,816 experimental datasets. The resulting HRAlit represents 23 tables with 21,704,001 records including 7 junction tables with 13,042,188 relationships. We demonstrate how HRAlit can be mined to identify leading experts, major papers, funding trends, or alignment with existing ontologies in support of systematic HRA construction and usage. Data and code are at https://github.com/cns-iu/hra-literature.

### Table S1. HRAlit data dictionary.

**Panel A.** Table description.

| Table Name | Attribute | Description | | |
|---|---|---|---|---|
| hralit_author | Purpose | To store information about authors who have contributed to publications, as well as creators and reviewers involved in the Human Reference Atlas (HRA) effort. | | |
| | Fields | Column | Data type | Description |
| | | orcid | varchar | Unique identifier for each author, based on the ORCID ID system. |
| | | first_name | varchar | Author's first name. |
| | | last_name | varchar | Author's last name. |
| | | first_pubyear | int8 | The year in which the author's first publication appeared. |
| | | career_age | int8 | The length of the author's career in years. |
| | | involved_funding | int8 | An integer indicating the number of funding opportunities the author has been involved in. |
| | Relationships | Linked to "*hralit_publication_author*" table via "*orcid*".<br>Linked to "*hralit_author_institution*" table via "*orcid*". | | |
| | Notes | The "*orcid*" field is based on the ORCID ID system, which provides a persistent digital identifier for researchers.<br>The "*career_age*" field is the difference of the current year (2023) and first_pubyear. This field will need to be updated annually to correctly represent the length of the author's career.<br>The "*involved_funding*" field indicates the number of funded projects that have supported the author's publications as of August 2023. This field will need to be updated as the author secures new funding in the future. | | |

| | | | | |
|---|---|---|---|---|
| hralit_publication | Purpose | To store detailed information about individual publications. | | |
| | Fields | Column | Data type | Description |
| | | pmid | varchar | Unique identifier for each publication based on the PubMed ID. |
| | | doi | varchar | Digital Object Identifier for the publication. |
| | | pubyear | int8 | The year in which the publication appeared. |
| | | article_title | varchar | Title of the article. |
| | | journal_title | varchar | Journal in which the article was published. |
| | Relationships | Linked to "*hralit_publication_author*" table via "*pmid*". | | |
| | | Linked to "hralit_pub_funding_funder" via "*pmid*". | | |
| | Notes | The "*pmid*" field is a unique identifier based on the PubMed system, commonly used in biomedical literature. | | |
| | | The "*doi*" field is another unique identifier that provides a permanent link to the article. | | |
| hralit_institution | Purpose | To store detailed information about research institutions. | | |
| | Fields | Column | Data type | Description |
| | | soa_institution_id | varchar | Unique identifier for each institution, based on SemOpenAlex dataset. |
| | | ror | varchar | Research Organization Registry (ROR) identifier, a globally unique identifier for research organizations. |
| | | institution_name | varchar | Name of the institution. |
| | | institution_type | varchar | Type of institution (e.g., education, government, company, etc.) |
| | | country_code | varchar | ISO country code indicating the country where the institution is located. |
| | Relationships | Linked to "*hralit_author_institution*" table via "*soa_institution_id*". | | |
| | Notes | The SemOpenAlex portal is available at https://semopenalex.org. | | |
| | | The Research Organization Registry (ROR) portal can be found at https://ror.org. | | |
| hralit_author_institution | Purpose | To establish relationships between authors and the institutions with which they are affiliated. | | |
| | Fields | Column | Data type | Description |
| | | orcid | varchar | Unique identifier for each author, based on the ORCID ID system. |
| | | soa_institution_id | varchar | Unique identifier for each institution, based on SemOpenAlex dataset. |
| | Relationships | Linked to "*hralit_author*" table via "*orcid*". | | |
| | | Linked to "*hralit_institution* " table via "*soa_institution_id*". | | |
| | Notes | The Research Organization Registry (ROR) portal can be found at https://ror.org. | | |
| hralit_funding | Purpose | To store information about various funding sources that support research. | | |
| | Fields | Column | Data type | Description |
| | | funding_id | varchar | Unique identifier for each funding source. |
| | | acronym | varchar | Acronym or short name for the funding source. |
| | Relationships | Linked to "*hralit_pub_funding_funder*" table via "*funding_id*". | | |
| | Notes | The "*acronym*" field can be used for easier identification and referencing of funding sources. | | |
| hralit_funder_cleaned | Purpose | To store cleaned and standardized information about entities that fund research. | | |
| | Fields | Column | Data type | Description |
| | | soa_funder_id | varchar | Unique identifier for each funder, based on SemOpenAlex dataset. |
| | | funder_name | varchar | Full name of the funder. |
| | | country_code | varchar | ISO country code indicating the country where the funder is |

| Table | Section | Content | | |
|---|---|---|---|---|
| | | | | based. |
| | Relationships | Linked to "*hralit_pub_funding_funder*" table via "*soa_funder_id*". | | |
| | Notes | The data for this table comes from SemOpenAlex. | | |
| hralit_pub_funding_funder | Purpose | To link publications with their respective funding sources and funders. | | |
| | Fields | Column | Data type | Description |
| | | pmid | varchar | Unique identifier for each publication based on the PubMed ID. |
| | | funding_id | varchar | Unique identifier for each funding source. |
| | | acronym | varchar | Acronym or short name for the funding source. |
| | | funder_name_pubmed | varchar | Name of the funder as it appears in PubMed. |
| | | soa_funder_id | varchar | Unique identifier for each funder, based on SemOpenAlex dataset. |
| | | country | varchar | Country where the funder is based. |
| | Relationships | Linked to "*hralit_publication*" table via "*pmid*".<br>Linked to "*hralit_funding*" table via "*funding_id*".<br>Linked to "*hralit_funder_cleaned*" table via "*soa_funder_id*". | | |
| | Notes | This table serves as a junction table to establish many-to-many relationships among publications, fundings, and funders.<br>This table includes uncleaned funder data from PubMed and cleaned funder data from SemOpenAlex. | | |
| hralit_digital_objects | Purpose | To store information about various digital objects related to HRA. | | |
| | Fields | Column | Data type | Description |
| | | type | varchar | Type of digital object (e.g., asct-b, omap, 2d-ftu, vascular-geometry, ref-organ). |
| | | name | varchar | Name of the digital object. |
| | | version | varchar | Version of the digital object. |
| | | title | varchar | Title associated with the digital object. |
| | | license | varchar | License under which the digital object is released. |
| | | publisher | varchar | Entity that published the digital object. |
| | | hubmap_id | varchar | Unique identifier within the HuBMAP (Human BioMolecular Atlas Program). |
| | | doi | varchar | Digital Object Identifier for the object. |
| | Relationships | Linked to "*hralit_creator*" table via "*hubmap_id*".<br>Linked to "*hralit_reviewer*" table via "*hubmap_id*". | | |
| | Notes | The data for this HRA 5th release table is available at https://hubmapconsortium.github.io/ccf-releases/v1.4/docs. | | |
| hralit_asct_publication | Purpose | To store information about general references and specific references in ASCT+B Tables. | | |
| | Fields | Column | Data type | Description |
| | | organ | varchar | Organ related to the entity being discussed in the publication. |
| | | id | varchar | Identifier for the publication. |
| | | doi | varchar | Digital Object Identifier for the publication. |
| | | notes | varchar | Additional notes or comments related to the publication. |
| | | type | varchar | Type of the publication (e.g., general_publication, reference). |
| | Relationships | Linked to "*hralit_publication*" table via "*doi*" field.<br>Linked to "*hralit_organ*" table via "organ" field. | | |
| | Notes | The data for this table is sourced from the 5th release of the Anatomical Structures, Cell Types, and Biomarker (ASCT+B) Tables. | | |
| hralit_other_refs | Purpose | To store information of publications associated with CZ CELLxGENE and CellMarker. | | |
| | Fields | Column | Data type | Description |

| | | | | |
|---|---|---|---|---|
| | | pmid | varchar | Unique identifier for each publication based on the PubMed ID. |
| | | doi | varchar | Digital Object Identifier for the publication. |
| | | source | varchar | Source from which the publication information was obtained (i.e., cxg, cellmarker). |
| | Relationships | Linked to "*hralit_publication*" table via "*pmid*" field. | | |
| | Notes | The data for this table is sourced from CZ CELLxGene API and CellMarker. CellMarker data is available at http://xteam.xbio.top/CellMarker/download/Human_cell_markers.txt | | |
| hralit_creator | Purpose | To store information about the creators of various digital objects for HRA. | | |
| | Fields | Column | Data type | Description |
| | | organ | varchar | Organ related to the entity being discussed in the publication. |
| | | orcid | varchar | Unique identifier for each creator, based on the ORCID ID system. |
| | | full_name | varchar | Full name of the creator. |
| | | first_name | varchar | First name of the creator. |
| | | last_name | varchar | Last name of the creator. |
| | | version | varchar | Version of digital objects related to the creator's contribution. |
| | | name | varchar | Name of digital objects related to the creator's contribution. |
| | | type | varchar | Type of digital objects related to the creator's contribution. |
| | | organ | varchar | Organ related to the digital object that the creator created. |
| | Relationships | Linked to "*hralit_author*" table via "*orcid*" field.<br>Linked to "*hralit_digital_object_5th_release*" table via "*hubmap_id*" field. | | |
| | Notes | This table serves as a junction table to establish many-to-many relationships among creators, authors, and digital objects.<br>The data for this table is available at https://hubmapconsortium.github.io/ccf-releases/v1.4/docs. | | |
| hralit_reviewer | Purpose | To store information about the reviewers of various digital objects for HRA. | | |
| | Fields | Column | Data type | Description |
| | | organ | varchar | Organ related to the entity being discussed in the publication. |
| | | orcid | varchar | Unique identifier for each reviewer, based on the ORCID ID system. |
| | | full_name | varchar | Full name of the reviewer. |
| | | first_name | varchar | First name of the reviewer. |
| | | last_name | varchar | Last name of the reviewer. |
| | | version | varchar | Version of digital objects related to the reviewer's contribution. |
| | | name | varchar | Name of digital objects related to the reviewer's contribution. |
| | | type | varchar | Type of digital objects related to the reviewer's contribution. |
| | | organ | varchar | Organ related to the digital object that the creator created. |
| | Relationships | Linked to "*hralit_author*" table via "*orcid*" field.<br>Linked to "*hralit_digital_object_5th_release*" table via "*hubmap_id*" field. | | |
| | Notes | This table serves as a junction table to establish many-to-many relationships among reviewers, authors, digital objects.<br>The data for this table is available at https://hubmapconsortium.github.io/ccf-releases/v1.4/docs. | | |
| hralit_author_expertise | Purpose | To store information about the areas of expertise for each author. | | |
| | Fields | Column | Data type | Description |
| | | organ | varchar | Organ related to the entity being discussed in the publication. |
| | | orcid | varchar | Unique identifier for each creator, based on the ORCID ID system. |
| | | expertise | varchar | Description or name of the area of expertise. |

| | | | | |
|---|---|---|---|---|
| | Relationships | Linked to "*hralit_author*" table via "*orcid*" field. Linked to "*hralit_organ*" table via "*expertise*" field. | | |
| | Notes | The expertise is tagged based on the titles and keywords or MeSH terms author's publications. | | |
| hralit_anatomical_structures | Purpose | To store information about different anatomical structures. | | |
| | Fields | Column | Data type | Description |
| | | organ | varchar | Organ related to the entity being discussed in the publication. |
| | | id | varchar | Unique identifier for each anatomical structure, typically in the format of a prefixed namespace and numerical code (e.g., "UBERON:0000178", "FMA:62467", etc.). |
| | | rdfs_label | varchar | RDF Schema label for the anatomical structure. |
| | | name | varchar | Name of the anatomical structure. |
| | Relationships | Linked to "*hralit_triple*" table via "*id*" field. | | |
| | Notes | The data for this table is sourced from the 5th release of the Anatomical Structures, Cell Types, and Biomarker (ASCT+B) Tables. | | |
| hralit_cell_types | Purpose | To store information about different cell types. | | |
| | Fields | Column | Data type | Description |
| | | organ | varchar | Organ related to the entity being discussed in the publication. |
| | | id | varchar | Unique identifier for each cell type, typically in the format of a prefixed namespace and numerical code (e.g., "CL:0000945"). |
| | | rdfs_label | varchar | RDF Schema label for the cell type. |
| | | name | varchar | Name of the cell type. |
| | Relationships | Linked to "*hralit_triple*" table via "*id*" field. | | |
| | Notes | The data for this table is sourced from the 5th release of the Anatomical Structures, Cell Types, and Biomarker (ASCT+B) Tables. | | |
| hralit_biomarkers | Purpose | To store information about various biomarkers. | | |
| | Fields | Column | Data type | Description |
| | | organ | varchar | Organ related to the entity being discussed in the publication. |
| | | id | varchar | Unique identifier for each biomarker, typically in the format of a prefixed namespace and numerical code (e.g., "HGNC:1633"). |
| | | rdfs_label | varchar | RDF Schema label for the biomarker. |
| | | name | varchar | Name of the biomarker. |
| | | b_type | varchar | Type or category of the biomarker (e.g., gene, protein, etc.). |
| | Relationships | Linked to "*hralit_triple*" table via "*id*" field. | | |
| | Notes | The data for this table is sourced from the 5th release of the Anatomical Structures, Cell Types, and Biomarker (ASCT+B) Tables. | | |
| hralit_triple | Purpose | To store ontology-related data, potentially including relationships between various biomedical entities. | | |
| | Fields | Column | Data type | Description |
| | | row_id | varchar | Unique identifier for each row, typically in the format of a prefixed namespace and numerical code (e.g., "blood_11", etc.). |
| | | id | varchar | Identifier for the entity. |
| | | rdfs_label | varchar | RDF Schema label for the entity. |
| | | name | varchar | Name of the entity. |
| | | b_type | varchar | Type or category of the entity. |
| | | organ | varchar | Organ associated with the entity. |
| | | ontology_type | varchar | Type or category of the entity (i.e., AS, CT, B). |

| | | | | |
|---|---|---|---|---|
| | Relationships | Linked to "*hralit_anatomical_structures*" table via "*id*". | | |
| | | Linked to "*hralit_cell_types*" table via "*id*". | | |
| | | Linked to "*hralit_biomarkers*" table via "*id*". | | |
| | Notes | This table serves as a junction table to establish many-to-many relationships among anatomical_structure entities, cell type entities, and biomarker entities. | | |
| | | The data for this table is sourced from the 5th release of the Anatomical Structures, Cell Types, and Biomarker (ASCT+B) Tables. | | |
| hralit_donor | Purpose | To store comprehensive information about donors, including both biometric data and medical history. | | |
| | Fields | Column | Data type | Description |
| | | organ | varchar | Organ related to the entity being discussed in the publication. |
| | | donor_id | varchar | Unique identifier for each donor. |
| | | sex | varchar | Biological sex of the donor. |
| | | age | varchar | Age of the donor. |
| | | death_event | varchar | Event or circumstance leading to the donor's death. |
| | | source | varchar | Source from which the donor information was obtained (i.e., hubmap, cxg, gtex). |
| | | age_unit | varchar | Unit in which the age is measured (e.g., years). |
| | | weight | varchar | Weight of the donor. |
| | | weight_unit | varchar | Unit in which the weight is measured (e.g., kg, pounds). |
| | | height | varchar | Height of the donor. |
| | | height_unit | varchar | Unit in which the height is measured (e.g., cm, inches). |
| | | race | varchar | Race of the donor. |
| | | body_mass_index | varchar | Body Mass Index (BMI) of the donor. |
| | | body_mass_index_unit | varchar | Unit in which the BMI is measured. |
| | | blood_type | varchar | Blood type of the donor (e.g., A, B, AB, O). |
| | | rh_blood_group | varchar | Rhesus (Rh) blood group of the donor (e.g., Rh negative). |
| | | rh_factor | varchar | Rhesus (Rh) factor (e.g., positive or negative). |
| | | kidney_donor_profile_index | varchar | Kidney Donor Profile Index (KDPI) of the donor. |
| | | kidney_donor_profile_index_unit | varchar | Unit in which the KDPI is measured. |
| | | cause_of_death | varchar | Cause of the donor's death. |
| | | medical_history | varchar | Medical history of the donor. |
| | | mechanism_of_injury | varchar | Mechanism of any injury sustained by the donor. |
| | | social_history | varchar | Social history of the donor. |
| | | sex_ontology | varchar | Ontological classification of sex. |
| | | race_ontology | varchar | Ontological classification of race. |
| | Relationships | Linked to "*hralit_datasets*" table via "*donor_id*". | | |
| | Notes | The data for this table is sourced from the Human BioMolecular Atlas Program (HuBMAP) portal, CZ CELLxGENE portal, and Genotype-Tissue Expression (GTEx) portal. | | |
| hralit_dataset | Purpose | To store detailed information about datasets, including associated donor and sample details, metadata, and dataset status. | | |
| | Fields | Column | Data type | Description |
| | | organ | varchar | Organ related to the entity being discussed in the publication. |
| | | dataset_id | varchar | Unique identifier for each dataset. |
| | | organ_gtex_id | varchar | GTEx identifier for the organ related to the dataset. |
| | | donor_id | varchar | Identifier for the donor associated with the dataset. |
| | | individual_id | varchar | Identifier for the individual from whom the sample was taken. |
| | | protocols_used | varchar | Protocols used in the generation or analysis of the dataset. |

| | | |
|---|---|---|
| rin_score_from_paxgene | varchar | RIN score obtained from the PAXgene process. |
| rin_score_from_frozen | varchar | RIN score obtained from the frozen sample. |
| organ | varchar | Organ from which the sample was taken. |
| autolysis_score | varchar | Score indicating the level of tissue autolysis. |
| sample_ischemic_time | varchar | Time for which the sample underwent ischemia. |
| sample_type | varchar | Type of sample (e.g., normal). |
| pathology_notes | varchar | Notes related to pathology findings. |
| source | varchar | Source from which the dataset was obtained (i.e., hubmap, cxg, gtex). |
| dataset_hubmap_id | varchar | HuBMAP identifier for the dataset. |
| dataset_status | varchar | Current status of the dataset (e.g., published). |
| dataset_group_name | varchar | Name of the group responsible for the dataset. |
| dataset_group_uuid | varchar | UUID of the dataset group. |
| dataset_date_time_created | varchar | Date and time the dataset was created. |
| dataset_created_by_email | varchar | Email of the individual who created the dataset. |
| dataset_date_time_modified | varchar | Date and time the dataset was last modified. |
| dataset_modified_by_email | varchar | Email of the individual who last modified the dataset. |
| lab_id_or_name | varchar | Identifier or name of the lab responsible for the dataset. |
| dataset_data_types | varchar | Types of data included in the dataset. |
| dataset_portal_url | varchar | URL to access the dataset in a portal. |
| first_sample_hubmap_id | varchar | HubMAP identifier for the first sample. |
| first_sample_submission_id | varchar | Submission identifier for the first sample. |
| first_sample_uuid | varchar | UUID for the first sample. |
| first_sample_type | varchar | Type of the first sample. |
| first_sample_portal_url | varchar | Portal URL for the first sample. |
| organ_hubmap_id | varchar | HuBMAP identifier for the organ. |
| organ_submission_id | varchar | Submission identifier for the organ. |
| organ_uuid | varchar | UUID for the organ. |
| donor_submission_id | varchar | Submission identifier for the donor. |
| donor_uuid | varchar | UUID for the donor. |
| donor_group_name | varchar | Name of the donor group. |
| rui_location_hubmap_id | varchar | HuBMAP identifier for the RUI location. |
| rui_location_submission_id | varchar | Submission identifier for the RUI location. |
| rui_location_uuid | varchar | UUID for the RUI location. |
| sample_metadata_hubmap_id | varchar | HuBMAP identifier for sample metadata. |
| sample_metadata | varchar | Submission identifier for sample metadata. |

| | | | | |
|---|---|---|---|---|
| | | _submission_id | | |
| | | sample_metadata_uuid | varchar | UUID for sample metadata. |
| | | processed_dataset_uuid | varchar | UUID for the processed dataset. |
| | | processed_dataset_hubmap_id | varchar | HuBMAP identifier for the processed dataset. |
| | | processed_dataset_status | varchar | Status of the processed dataset. |
| | | processed_dataset_portal_url | varchar | Portal URL for the processed dataset. |
| | | previous_version_hubmap_ids | varchar | HuBMAP identifiers for previous versions of the dataset. |
| | | cxg_dataset_id | varchar | Identifier for the CxG dataset. |
| | | dataset_title | varchar | Title of the dataset. |
| | | dataset_h5ad_path | varchar | Path to the H5AD file for the dataset. |
| | | dataset_total_cell_count | varchar | Total cell count in the dataset. |
| | | collection_id | varchar | Identifier for the collection to which the dataset belongs. |
| | | collection_name | varchar | Name of the collection. |
| | | publication_doi | varchar | DOI for publications related to the dataset. |
| | | organ_ontology | varchar | Ontological classification for the organ. |
| | | anatomical_structure | varchar | Description of the anatomical structure from which the sample was taken. |
| | | anatomical_structure_ontology | varchar | Ontological classification for the anatomical structure. |
| | | suspension_type | varchar | Type of suspension used in the sample. |
| | Relationships | Linked to "*hralit_donor*" table via "*donor_id*".<br>Linked to "*hralit_publication*" table via "*publication_doi*". | | |
| | Notes | The data for this table is sourced from the Human BioMolecular Atlas Program (HuBMAP) portal, CZ CELLxGENE portal, and Genotype-Tissue Expression (GTEx) portal.<br>This table serves as a junction table to establish many-to-many relationships among datasets, donors, publications. | | |
| hralit_publication_subject | Purpose | To store information about the organ subjects that are the focus of various publications. | | |
| | Fields | **Column** | **Data type** | **Description** |
| | | organ | varchar | Organ related to the entity being discussed in the publication. |
| | | pmid | varchar | Unique identifier for each publication based on the PubMed ID. |
| | | organ | varchar | The subject of the publication. |
| | Relationships | Linked to the "*hralit_publication*" table via "*pmid*" field. | | |
| | Notes | The organ subject is tagged based on the titles and keywords or MeSH terms of the publication. | | |
| hralit_publication_author | Purpose | To establish relationships between publications and their respective authors | | |
| | Fields | **Column** | **Data type** | **Description** |
| | | pmid | varchar | Unique identifier for each publication based on the PubMed ID. |
| | | orcid | varchar | Unique identifier for each author, based on the ORCID ID system. |
| | Relationships | Linked to "*hralit_publication*" table via "*pmid*".<br>Linked to "*hralit_author*" table via "*orcid*". | | |
| | Notes | This table serves as a junction table to establish many-to-many relationships between publications and authors. | | |

| | | | | |
|---|---|---|---|---|
| hralit_organ | Purpose | To store information about organs in 5th release ASCT+B Tables. | | |
| | Fields | Column | Data type | Description |
| | | organ | varchar | Organ name. |
| | Relationships | Linked to "*hralit_triple*" table via "*organ*". | | |
| | | Linked to "*hralit_publication_subject*" table via "*organ*". | | |
| | | Linked to "*hralit_asctb_publication*" table via "*organ*". | | |
| | | Linked to "*hralit_author_expertise*" table via "*orcid*". | | |
| | Notes | The data for this table is available at https://hubmapconsortium.github.io/ccf-releases/v1.4/docs. | | |
| hralit_asctb_linkage | Purpose | To store ontology-related data, potentially including relationships between various biomedical entities. | | |
| | Fields | Column | Data type | Description |
| | | id | varchar | Identifier for the entity. |
| | | source_id | varchar | Identifier for the entity. |
| | | source_rdfs_label | varchar | RDF Schema label for the entity. |
| | | source_name | varchar | Name of the entity. |
| | | source_type | varchar | Type or category of the entity. |
| | | relationship | varchar | Linkage between source and target. |
| | | target_id | varchar | Identifier for the entity. |
| | | target_rdfs_label | varchar | RDF Schema label for the entity. |
| | | target_name | varchar | Name of the entity. |
| | | target_type | varchar | Type or category of the entity. |
| | Relationships | Linked to "*hralit_anatomical_structures*" table via "*id*". | | |
| | | Linked to "*hralit_cell_types*" table via "*id*". | | |
| | | Linked to "*hralit_biomarkers*" table via "*id*". | | |
| | Notes | This table serves as a junction table to establish many-to-many relationships among anatomical_structure entities, cell type entities, and biomarker entities. | | |
| | | The data for this table is sourced from the 5th release of the Anatomical Structures, Cell Types, and Biomarker (ASCT+B) Tables. | | |

**Panel B.** Row count statistics for 23 HRAlit database tables.

| Table | Rows | Columns |
|---|---:|---:|
| hralit_anatomical_structures | 4,383 | 3 |
| hralit_asctb_publication | 1,288 | 5 |
| hralit_asctb_linkage | 20,146 | 9 |
| hralit_author | 583,117 | 6 |
| hralit_author_expertise | 768,178 | 2 |
| hralit_author_institution | 464,043 | 2 |
| hralit_biomarkers | 3,234 | 4 |
| hralit_cell_types | 1,422 | 3 |
| hralit_creator | 550 | 9 |
| hralit_dataset | 7,337 | 52 |
| hralit_digital_objects | 295 | 8 |
| hralit_donor | 4,639 | 24 |
| hralit_funder_cleaned | 6,427 | 3 |
| hralit_funding | 917,061 | 2 |
| hralit_institution | 26,235 | 5 |
| hralit_organ | 31 | 1 |
| hralit_other_refs | 1,823 | 3 |
| hralit_pub_funding_funder | 2,632,888 | 6 |
| hralit_publication | 7,103,369 | 5 |

| | | |
|---|---:|---:|
| hralit_publication_author | 1,079,698 | 2 |
| hralit_publication_subject | 7,898,258 | 2 |
| hralit_reviewer | 602 | 9 |
| hralit_triple | 178,977 | 7 |
| **Total** | **21,704,001** | **172** |

Note that hralit_asctb_linkage, hralit_author_expertise, hralit_author_institution, hralit_pub_funding_funder, hralit_publication_author, hralit_publication_subject, hralit_triple are the 7 junction tables.

**Panel C.** Node count statistics for HRAlit database tables.

| Name | #Nodes |
|---|---:|
| anatomical structures | 2,689 |
| asctb publications | 1,057 |
| authors | 583,117 |
| biomarkers | 1,770 |
| cell types | 590 |
| CellMarker/CxG/GTEx publications* | 1,816 |
| creators | 101 |
| datasets | 1,816 |
| digital objects | 295 |
| donors | 4,639 |
| funders (cleaned) | 6,427 |
| funders (uncleaned) | 63,691 |
| funded projects | 896,680 |
| institutions (cleaned) | 26,235 |
| organs | 31 |
| publications | 7,103,180 |
| reviewers | 99 |
| **Total** | **8,694,233** |

Note that:
In CellMarker, CxG, and GTEx publications, the number refers to the publications with DOIs.
The number of anatomical structures refers to the number of anatomical structures that have ontology IDs.
The number of cell types refers to the number of cell types that have ontology IDs.
The number of biomarkers refers to the number of biomarkers that have ontology IDs.

**Panel D.** Linkage count statistics for HRAlit database for 22 relationship types.

| Relationships | #Linkages |
|---|---:|
| author & institution | 464,043 |
| AS & CT &B | 178,977 |
| publication & organ | 7,898,258 |
| asctb_refs & publication | 472 |
| asctb_refs & organ | 1,288 |
| creator & author | 550 |
| reviewer & author | 602 |
| dataset & donor | 7,337 |
| dataset & publication | 5,228 |
| creator & digital object | 550 |
| reviewer & digital object | 602 |
| author & organ | 768,178 |
| publication & funding | 2,616,446 |
| funding & cleaned funder | 147,202 |
| funding & uncleaned funder | 888,249 |
| triple & organ | 17,092 |
| CellMarker/CxG refs & publication | 1,817 |

| | |
|---|---|
| publication & author | 1,079,698 |
| AS & AS | 4,835 |
| AS & CT | 7,879 |
| CT & CT | 620 |
| CT & B | 6,812 |
| **Total** | **14,096,735** |

### Table S2. Expert, Literature, and Experimental Data Evidence for HRA.

**Panel A.** Overview.

| Organ | #Datasets | #Cells | #Experts | Avg. *h*-index | #Fundings | #Funders | #Publications | Total citations |
|---|---|---|---|---|---|---|---|---|
| blood pelvis | 1,788 | 1,952,941,521 | 3,087 | 0.6718 | 2237 | 131 | 39,382 | 460,862 |
| blood vasculature | 17 | 3,311,718 | 79 | 1.0000 | 335 | 17 | 326 | 8657 |
| bone marrow | 24 | 4,664,385 | 14,625 | 0.9442 | 45667 | 566 | 174,334 | 5,252,011 |
| brain | 1,666 | 244,852,929 | 122,304 | 0.7521 | 251,714 | 2,703 | 1,129,073 | 36,301,681 |
| eye | 256 | 14,844,890 | 25,131 | 0.7882 | 46,232 | 806 | 306,681 | 5,336,058 |
| fallopian tube | 69 | 3,519,078 | 1,055 | 0.9223 | 1,768 | 71 | 18,142 | 266,846 |
| heart | 49 | 4,769,948 | 76,545 | 0.5986 | 117,341 | 1,583 | 1,019,527 | 19,501,222 |
| kidney | 625 | 19,329,269 | 59,910 | 0.7027 | 97,041 | 1,485 | 762,095 | 15,329,887 |
| knee | 6 | 0 | 17,168 | 0.4733 | 12,381 | 602 | 141,568 | 2,478,738 |
| large intestine | 289 | 7,823,252 | 405 | 1.2444 | 955 | 29 | 10,531 | 204,088 |
| liver | 89 | 9,448,009 | 152,354 | 0.7295 | 192,239 | 2,911 | 1,288,368 | 28,725,600 |
| lung | 506 | 415,050,060 | 79,730 | 0.6982 | 119,090 | 1,864 | 688,820 | 14,009,431 |
| lymph node | 55 | 4,295,177 | 16,076 | 0.7837 | 24,443 | 509 | 154,950 | 3,325,388 |
| lymph vasculature | 0 | 0 | 70 | 0.9286 | 236 | 11 | 214 | 8,885 |
| main bronchus | 0 | 0 | 31 | 0.1935 | 18 | 3 | 668 | 2,044 |
| muscular system | 37 | 7,618,410 | 633 | 0.8167 | 549 | 45 | 2,029 | 31,782 |
| ovary | 6 | 156,804 | 7,716 | 0.7381 | 16,950 | 417 | 93,764 | 2,251,351 |
| pancreas | 46 | 4,539,924 | 7,124 | 0.7887 | 14,925 | 316 | 99,427 | 2,171,132 |
| peripheral nervous system | 0 | 0 | 2,908 | 0.9646 | 7667 | 162 | 34,753 | 835,163 |
| placenta | 0 | 0 | 9,971 | 0.7797 | 17,688 | 468 | 86,872 | 1,934,916 |
| prostate | 73 | 4,159,866 | 23,131 | 0.6877 | 34,219 | 907 | 174,800 | 3,671,783 |
| skeletal | 0 | 0 | 40,324 | 0.8397 | 80,013 | 1,243 | 284,538 | 8,581,991 |
| skin | 58 | 5,066,152 | 56,735 | 0.7685 | 75,199 | 1,421 | 611,054 | 11,687,148 |
| small intestine | 308 | 7,153,510 | 3,796 | 0.8725 | 9,850 | 214 | 68,140 | 1,498,261 |
| spinal cord | 134 | 7,280,191 | 14,357 | 0.7689 | 30,421 | 656 | 161,645 | 4,159,877 |
| spleen | 68 | 3,039,279 | 5,795 | 1.1757 | 30,476 | 338 | 125,280 | 3,484,733 |
| thymus | 25 | 0 | 3,653 | 0.9296 | 14,510 | 154 | 62,054 | 2,082,347 |
| trachea | 0 | 0 | 5,935 | 0.6056 | 9,532 | 199 | 91,543 | 1,281,223 |
| ureter | 1 | 598,266 | 3,921 | 0.5376 | 3,294 | 144 | 62,702 | 593,136 |
| urinary bladder | 35 | 4,767,426 | 10,343 | 0.7315 | 14,713 | 460 | 133,489 | 2,082,534 |

| | | | | | | | | |
|---|---|---|---|---|---|---|---|---|
| uterus | 239 | 4,657,715 | 3,266 | 0.7676 | 8,470 | 177 | 71,489 | 1,258,343 |

**Panel B.** Average citations per organ.

| Organ | Avg. Citation | Organ | Avg. Citation |
|---|---|---|---|
| blood pelvis | 11.7024 | ovary | 24.0108 |
| blood vasculature | 26.5552 | pancreas | 21.8364 |
| bone marrow | 30.1261 | peripheral nervous system | 24.0314 |
| brain | 32.1518 | placenta | 22.2732 |
| eye | 17.3994 | prostate | 21.0056 |
| fallopian tube | 14.7087 | skeletal | 30.1611 |
| heart | 19.1277 | skin | 19.1262 |
| kidney | 20.1155 | small intestine | 21.9880 |
| knee | 17.5092 | spinal cord | 25.7346 |
| large intestine | 19.3797 | spleen | 27.8156 |
| liver | 22.2961 | thymus | 33.5570 |
| lung | 20.3383 | trachea | 13.9959 |
| lymph node | 21.4610 | ureter | 9.4596 |
| lymph vasculature | 41.5187 | urinary bladder | 15.6008 |
| main bronchus | 3.0599 | uterus | 17.6019 |
| muscular system | 15.6639 | | |

**Panel C.** Total amount in U.S. dollars for six additional funders.

| Organ | ARDC | CIHR | EC |
|---|---|---|---|
| blood pelvis | $6,473,386.23 | $0.00 | $4,495,388.82 |
| blood vasculature | $1,947,038,411.00 | $0.00 | $2,864,292.30 |
| bone marrow | $763,369,441.00 | $13,259,843.05 | $209,817,035.80 |
| brain | $1,873,361,481.00 | $137,575,676.40 | $4,145,944,813.00 |
| eye | $343,188,660.90 | $20,345,718.40 | $1,053,460,297.00 |
| fallopian tube | $317,562,288.50 | $0.00 | $5,700,042.76 |
| heart | $1,244,839,814.00 | $160,900,846.30 | $2,199,690,627.00 |
| kidney | $481,300,595.30 | $22,003,833.66 | $566,944,165.20 |
| knee | $96,506,817.78 | $4,005,468.12 | $109,795,367.50 |
| large intestine | $2,202,622,468.00 | $0.00 | $4,646,794.04 |
| liver | $278,793,954.60 | $128,564,824.40 | $18,003,210,379.00 |
| lung | $590,666,713.40 | $57,502,762.27 | $708,065,189.80 |
| lymph node | $345,818,098.90 | $1,254,000.05 | $55,501,908.86 |
| lymph vasculature | $75,573,224.31 | $0.00 | $0.00 |
| main bronchus | $285,636,866.20 | $0.00 | $0.00 |
| muscular system | $4,765,955,702.00 | $0.00 | $21,871,483.24 |
| ovary | $104,570,502.10 | $3,667,175.69 | $36,773,882.17 |
| pancreas | $55,424,017.84 | $6,475,765.91 | $134,472,313.50 |
| peripheral nervous system | $4,106,268,238.00 | $0.00 | $17,977,376.77 |
| placenta | $73,898,069.20 | $25,484,893.29 | $152,096,551.40 |
| prostate | $212,314,077.20 | $16,798,601.78 | $317,843,336.40 |
| skeletal | $189,137,621.10 | $120,703,280.20 | $590,682,929.00 |
| skin | $355,154,230.90 | $19,621,283.14 | $1,252,143,512.00 |
| small intestine | $1,271,367,551.00 | $389,006.29 | $9,004,079.74 |
| spinal cord | $450,669,418.70 | $46,699,649.68 | $200,696,263.50 |
| spleen | $7,079,231.22 | $1,208,493.90 | $35,367,381.96 |
| thymus | $21,596,410.36 | $2,477,524.62 | $48,758,008.47 |
| trachea | $563,718.77 | $0.00 | $13,496,707.47 |

| | | | |
|---|---|---|---|
| ureter | $544,029.89 | $151,741.60 | $173,753.03 |
| urinary bladder | $328,413,111.70 | $0.00 | $0.00 |
| uterus | $68,744,457.70 | $901,968.19 | $19,304,420.24 |

| Organ | KAKEN | NIH | NSF |
|---|---|---|---|
| blood pelvis | $21,581,073.24 | $988,938,755.00 | $0.00 |
| blood vasculature | $478,637.95 | $4,585,407.00 | $0.00 |
| bone marrow | $874,646,679.50 | $14,137,363,749.00 | $0.00 |
| brain | $4,309,529,730.00 | $94,719,648,826.00 | $0.00 |
| eye | $1,023,130,174.00 | $15,251,379,619.00 | $0.00 |
| fallopian tube | $25,307,627.84 | $103,348,182.00 | $0.00 |
| heart | $1,505,035,718.00 | $48,021,255,774.00 | $0.00 |
| kidney | $1,052,887,557.00 | $26,025,180,482.00 | $0.00 |
| knee | $196,981,780.20 | $1,906,649,921.00 | $0.00 |
| large intestine | $59,680,068.28 | $721,265,466.00 | $0.00 |
| liver | $2,093,888,258.00 | $54,487,038,268.00 | $0.00 |
| lung | $1,582,072,958.00 | $47,370,697,019.00 | $0.00 |
| lymph node | $410,770,390.10 | $6,534,733,532.00 | $0.00 |
| lymph vasculature | $28,067,088.92 | $0.00 | $0.00 |
| main bronchus | $3,342,252.59 | $4,054,737.00 | $16,904,983,552.00 |
| muscular system | $203,980,245.70 | $371,709,982.00 | $91,682,007,165.00 |
| ovary | $331,769,060.30 | $6,529,175,563.00 | $0.00 |
| pancreas | $364,198,138.80 | $9,281,635,175.00 | $0.00 |
| peripheral nervous system | $36,414,355.24 | $2,335,561,740.00 | $0.00 |
| placenta | $201,593,755.70 | $3,640,228,212.00 | $0.00 |
| prostate | $522,814,503.40 | $12,732,528,277.00 | $0.00 |
| skeletal | $1,028,129,453.00 | $16,302,837,412.00 | $0.00 |
| skin | $1,289,075,495.00 | $18,263,992,502.00 | $0.00 |
| small intestine | $213,412,561.60 | $1,729,262,965.00 | $0.00 |
| spinal cord | $529,243,963.20 | $6,103,153,284.00 | $0.00 |
| spleen | $191,996,715.00 | $2,999,216,626.00 | $0.00 |
| thymus | $340,890,188.40 | $6,464,796,621.00 | $0.00 |
| trachea | $36,951,997.83 | $835,885,868.00 | $18,454,065.00 |
| ureter | $10,812,928.57 | $429,161,425.00 | $0.00 |
| urinary bladder | $87,454,286.13 | $1,903,866,962.00 | $0.00 |
| uterus | $156,536,198.70 | $1,916,656,866.00 | $32,225,240.00 |

**Panel D.** The number of publications per publication year.

| #Pubmed Publications | | | | | | | | | | | | | |
|---|---|---|---|---|---|---|---|---|---|---|---|---|---|
| Year | #P | Year | #P | Year | #P | Year | #P | Year | #P | Year | #P | Year | #P |
| 1781 | 1 | 1820 | 19 | 1854 | 48 | 1888 | 131 | 1922 | 204 | 1956 | 16,849 | 1990 | 87,347 |
| 1785 | 4 | 1821 | 23 | 1855 | 69 | 1889 | 159 | 1923 | 258 | 1957 | 17,651 | 1991 | 86,109 |
| 1786 | 6 | 1822 | 19 | 1856 | 73 | 1890 | 136 | 1924 | 259 | 1958 | 17,285 | 1992 | 86,745 |
| 1787 | 3 | 1823 | 23 | 1857 | 41 | 1891 | 137 | 1925 | 286 | 1959 | 18,221 | 1993 | 88,135 |
| 1788 | 3 | 1824 | 21 | 1858 | 78 | 1892 | 171 | 1926 | 303 | 1960 | 17,748 | 1994 | 91,534 |
| 1789 | 1 | 1825 | 15 | 1859 | 53 | 1893 | 172 | 1927 | 335 | 1961 | 19,986 | 1995 | 94,372 |
| 1790 | 7 | 1826 | 26 | 1860 | 70 | 1894 | 207 | 1928 | 275 | 1962 | 21,071 | 1996 | 95,364 |
| 1791 | 2 | 1827 | 19 | 1861 | 41 | 1895 | 182 | 1929 | 297 | 1963 | 27,886 | 1997 | 96,695 |
| 1792 | 3 | 1828 | 37 | 1862 | 36 | 1896 | 208 | 1930 | 335 | 1964 | 36,416 | 1998 | 99,450 |
| 1795 | 1 | 1829 | 43 | 1863 | 31 | 1897 | 214 | 1931 | 307 | 1965 | 32,677 | 1999 | 99,514 |
| 1796 | 1 | 1830 | 50 | 1864 | 33 | 1898 | 223 | 1932 | 314 | 1966 | 34,089 | 2000 | 104,895 |
| 1797 | 6 | 1831 | 29 | 1865 | 39 | 1899 | 193 | 1933 | 307 | 1967 | 39,932 | 2001 | 105,023 |
| 1798 | 2 | 1832 | 20 | 1866 | 82 | 1900 | 175 | 1934 | 282 | 1968 | 46,099 | 2002 | 107,670 |
| 1799 | 13 | 1833 | 23 | 1867 | 79 | 1901 | 212 | 1935 | 322 | 1969 | 50,133 | 2003 | 114,387 |
| 1800 | 19 | 1834 | 26 | 1868 | 63 | 1902 | 177 | 1936 | 309 | 1970 | 51,135 | 2004 | 119,323 |
| 1801 | 8 | 1835 | 31 | 1869 | 92 | 1903 | 219 | 1937 | 307 | 1971 | 55,340 | 2005 | 124,430 |

| | | | | | | | | | | | | | |
|---|---|---|---|---|---|---|---|---|---|---|---|---|---|
| 1802 | 9 | 1836 | 16 | 1870 | 76 | 1904 | 236 | 1938 | 303 | 1972 | 58,161 | 2006 | 129,535 |
| 1803 | 7 | 1837 | 15 | 1871 | 88 | 1905 | 202 | 1939 | 290 | 1973 | 59,613 | 2007 | 133,738 |
| 1804 | 10 | 1838 | 20 | 1872 | 141 | 1906 | 192 | 1940 | 231 | 1974 | 61,470 | 2008 | 139,676 |
| 1805 | 18 | 1839 | 16 | 1873 | 127 | 1907 | 230 | 1941 | 282 | 1975 | 60,401 | 2009 | 144,807 |
| 1806 | 10 | 1840 | 30 | 1874 | 136 | 1908 | 294 | 1942 | 241 | 1976 | 57,485 | 2010 | 151,875 |
| 1807 | 15 | 1841 | 86 | 1875 | 140 | 1909 | 321 | 1943 | 216 | 1977 | 58,798 | 2011 | 158,804 |
| 1808 | 6 | 1842 | 89 | 1876 | 125 | 1910 | 283 | 1944 | 190 | 1978 | 60,537 | 2012 | 172,154 |
| 1809 | 13 | 1843 | 100 | 1877 | 115 | 1911 | 290 | 1945 | 2,189 | 1979 | 63,994 | 2013 | 181,706 |
| 1810 | 12 | 1844 | 60 | 1878 | 183 | 1912 | 303 | 1946 | 6,893 | 1980 | 64,620 | 2014 | 188,495 |
| 1811 | 14 | 1845 | 59 | 1879 | 134 | 1913 | 274 | 1947 | 8,107 | 1981 | 65,212 | 2015 | 193,971 |
| 1812 | 10 | 1846 | 77 | 1880 | 120 | 1914 | 266 | 1948 | 9,017 | 1982 | 68,497 | 2016 | 193,013 |
| 1813 | 20 | 1847 | 53 | 1881 | 149 | 1915 | 220 | 1949 | 8,709 | 1983 | 71,438 | 2017 | 195,421 |
| 1814 | 24 | 1848 | 49 | 1882 | 120 | 1916 | 187 | 1950 | 12,155 | 1984 | 75,162 | 2018 | 202,447 |
| 1815 | 14 | 1849 | 74 | 1883 | 113 | 1917 | 199 | 1951 | 14,007 | 1985 | 77,104 | 2019 | 211,480 |
| 1816 | 29 | 1850 | 64 | 1884 | 119 | 1918 | 172 | 1952 | 15,271 | 1986 | 77,636 | 2020 | 228,935 |
| 1817 | 15 | 1851 | 55 | 1885 | 114 | 1919 | 174 | 1953 | 16,058 | 1987 | 78,949 | 2021 | 242,865 |
| 1818 | 20 | 1852 | 68 | 1886 | 146 | 1920 | 188 | 1954 | 16,548 | 1988 | 80,631 | 2022 | 234,436 |
| 1819 | 13 | 1853 | 49 | 1887 | 169 | 1921 | 235 | 1955 | 17,274 | 1989 | 84,882 | 2023 | 52,676 |

**#CZ CELLxGENE Publications**

| Year | #P | Year | #P | Year | #P | Year | #P | Year | #P | Year | #P |
|---|---|---|---|---|---|---|---|---|---|---|---|
| 2018 | 3 | 2019 | 4 | 2020 | 12 | 2021 | 14 | 2022 | 14 | 2023 | 2 |

**#CellMarker Publications**

| Year | #P | Year | #P | Year | #P | Year | #P | Year | #P | Year | #P | Year | #P |
|---|---|---|---|---|---|---|---|---|---|---|---|---|---|
| 1978 | 1 | 1985 | 2 | 1992 | 10 | 1999 | 21 | 2006 | 43 | 2013 | 151 | 2020 | |
| 1979 | 2 | 1986 | | 1993 | 3 | 2000 | 23 | 2007 | 39 | 2014 | 177 | 2021 | |
| 1980 | | 1987 | | 1994 | 8 | 2001 | 26 | 2008 | 54 | 2015 | 173 | 2022 | |
| 1981 | 1 | 1988 | 3 | 1995 | 11 | 2002 | 33 | 2009 | 81 | 2016 | 137 | 2023 | |
| 1982 | 1 | 1989 | 10 | 1996 | | 2003 | 24 | 2010 | 89 | 2017 | 154 | | |
| 1983 | 1 | 1990 | 19 | 1997 | 14 | 2004 | 34 | 2011 | 106 | 2018 | 84 | | |
| 1984 | 4 | 1991 | 10 | 1998 | 14 | 2005 | 32 | 2012 | 124 | 2019 | 3 | | |

**Panel E.** Correlation coefficient matrix of variables.

| | #Datasets | #Cells | #Experts | Avg. *h*-index | #Fundings | #Funders | #Publications | Total citations | Total award amount |
|---|---|---|---|---|---|---|---|---|---|
| #Datasets | 1 | | | | | | | | |
| #Cells | 0.77 | 1 | | | | | | | |
| #Experts | 0.332 | 0.014 | 1 | | | | | | |
| Avg. *h*-index | -0.058 | -0.119 | -0.175 | 1 | | | | | |
| #Fundings | 0.432 | 0.014 | 0.964 | -0.109 | 1 | | | | |
| #Funders | 0.332 | -0.003 | 0.983 | -0.176 | 0.967 | 1 | | | |
| #Publications | 0.329 | -0.003 | 0.976 | -0.186 | 0.949 | 0.961 | 1 | | |
| Total citations | 0.43 | 0.008 | 0.967 | -0.132 | 0.992 | 0.958 | 0.967 | 1 | |
| Total award amount | 0.003 | -0.096 | 0.036 | 0.254 | 0.027 | -0.026 | 0.009 | 0.038 | 1 |

### Table S3. HRAlit diversity.

**Panel A.** Gender in each age group.

| SURVEY | | |
|---|---|---|
| **Age Group** | **#Male** | **#Female** |
| 25 - 34 | 1 | 3 |
| 35 - 44 | 5 | 4 |
| 45 - 54 | 0 | 2 |
| 55 - 64 | 2 | 2 |
| 65 - 74 | 2 | 0 |
| 75 - 84 | 1 | 0 |
| **DONOR** | | |
| **Age Group** | **#Male** | **#Female** |
| 0-9 | 1 | 0 |
| 10-19 | 11 | 9 |
| 20-29 | 320 | 247 |

| | | |
|---|---|---|
| 30-39 | 230 | 325 |
| 40-49 | 353 | 339 |
| 50-59 | 414 | 324 |
| 60-69 | 342 | 407 |
| 70-79 | 341 | 275 |
| 80-89 | 219 | 365 |
| 90-99 | 7 | 7 |
| Unknown | | 103 |
| | **WORLD POPULATION** | |
| **Age Group** | **#Male** | **#Female** |
| 0-9 | 688,798,028 | 688,798,028 |
| 10-19 | 675,087,653 | 675,087,653 |
| 20-29 | 617,612,692 | 617,612,692 |
| 30-39 | 601,856,786 | 601,856,786 |
| 40-49 | 503,847,259 | 503,847,259 |
| 50-59 | 435,282,669 | 435,282,669 |
| 60-69 | 298,238,474 | 298,238,474 |
| 70-79 | 161,315,546 | 161,315,546 |
| 80-89 | 53,672,265 | 53,672,265 |
| 90-99 | 7,145,839 | 7,145,839 |
| 100+ | 130,480 | 130,480 |

**Panel B.** Gender in each career age.

| Career Age | #Male Authors | #Female Authors | #Male Experts | #Female Experts |
|---|---|---|---|---|
| 1 | 16 | 17 | 1 | 0 |
| 2 | 121 | 80 | 4 | 8 |
| 3 | 259 | 179 | 14 | 8 |
| 4 | 427 | 326 | 11 | 13 |
| 5 | 724 | 447 | 6 | 3 |
| 6 | 13,193 | 8,678 | 9 | 11 |
| 7 | 16,244 | 9,597 | 15 | 10 |
| 8 | 6,668 | 3,538 | 7 | 2 |
| 9 | 2,244 | 1,143 | 0 | 0 |
| 10 | 1,753 | 991 | 0 | 0 |
| 11 | 49 | 22 | 0 | 0 |
| 12 | 35 | 24 | 0 | 0 |
| 13 | 10 | 3 | 0 | 0 |
| 14 | 6 | 1 | 0 | 0 |
| 15 | 4 | 0 | 0 | 0 |
| 16 | 1 | 2 | 0 | 0 |
| 17 | 2 | 0 | 0 | 0 |

**Panel C.** Distribution of surveys, authors, experts, and donors by race.

| Race | #Survey | #Authors | #Experts | #Donors |
|---|---|---|---|---|
| Asian | 2 | 15,864 | 38 | 648 |
| Hispanic | 1 | 12720 | 13 | 54 |
| Black | 0 | 5827 | 10 | 107 |
| White | 17 | 22380 | 87 | 2834 |
| Other | 0 | 10012 | 12 | 52 |
| Unknown | 4 | 0 | 0 | 944 |

**Panel D.** Number of authors by country code.

| CC | #Authors | CC | #Authors | CC | #Authors | CC | #Authors | CC | #Authors |
|---|---:|---|---:|---|---:|---|---:|---|---:|
| AD | 2 | CV | 2 | IL | 2,432 | MT | 49 | SK | 438 |
| AE | 435 | CW | 4 | IM | 1 | MU | 11 | SL | 6 |
| AF | 11 | CY | 228 | IN | 8,722 | MV | 5 | SN | 20 |
| AG | 65 | CZ | 1,832 | IQ | 202 | MW | 53 | SO | 10 |
| AL | 15 | DE | 19,896 | IR | 5,050 | MX | 2,914 | SR | 6 |
| AM | 49 | DK | 4,698 | IS | 163 | MY | 1,521 | SS | 6 |
| AO | 4 | DO | 12 | IT | 22,408 | MZ | 20 | ST | 15 |
| AR | 1,096 | DZ | 75 | JE | 4 | NA | 16 | SV | 3 |
| AT | 2,933 | EC | 152 | JM | 24 | NC | 6 | SX | 4 |
| AU | 14,466 | EE | 151 | JO | 446 | NE | 8 | SY | 60 |
| AZ | 46 | EG | 2,792 | JP | 19,956 | NG | 571 | SZ | 2 |
| BA | 69 | ER | 5 | KE | 162 | NI | 8 | TC | 1 |
| BB | 13 | ES | 14,497 | KG | 27 | NL | 9,239 | TD | 1 |
| BD | 488 | ET | 778 | KH | 87 | NO | 2,523 | TG | 11 |
| BE | 3,952 | FI | 2,523 | KN | 21 | NP | 223 | TH | 1,869 |
| BF | 40 | FJ | 9 | KR | 16,259 | NZ | 1,434 | TJ | 30 |
| BG | 285 | FK | 1 | KW | 129 | OM | 108 | TN | 407 |
| BH | 44 | FO | 6 | KY | 2 | PA | 47 | TO | 2 |
| BI | 10 | FR | 12,112 | KZ | 166 | PE | 364 | TR | 9,385 |
| BJ | 10 | GA | 5 | LA | 3 | PF | 6 | TT | 32 |
| BM | 1 | GB | 25,287 | LB | 328 | PG | 5 | TW | 5,130 |
| BN | 22 | GD | 23 | LC | 3 | PH | 190 | TZ | 167 |
| BO | 19 | GE | 96 | LI | 2 | PK | 1,397 | UA | 331 |
| BR | 16,105 | GF | 6 | LK | 213 | PL | 7,299 | UG | 221 |
| BS | 2 | GH | 292 | LS | 1 | PS | 51 | US | 103,179 |
| BT | 6 | GI | 3 | LT | 322 | PT | 3,913 | UY | 147 |
| BW | 26 | GL | 5 | LU | 135 | PY | 23 | UZ | 22 |
| BY | 75 | GM | 19 | LV | 134 | QA | 407 | VC | 1 |
| CA | 12,751 | GN | 5 | LY | 12 | RE | 11 | VE | 36 |
| CD | 36 | GP | 11 | MA | 254 | RO | 1,578 | VG | 1 |
| CF | 1 | GR | 2,305 | MC | 16 | RS | 758 | VN | 423 |
| CG | 12 | GT | 22 | MD | 6 | RU | 5,458 | WS | 3 |
| CH | 5,831 | GW | 5 | ME | 7 | RW | 33 | XK | 16 |
| CI | 10 | GY | 1 | MG | 17 | SA | 2,335 | YE | 33 |
| CL | 1,066 | HN | 10 | MK | 45 | SB | 1 | ZA | 1,576 |
| CM | 112 | HR | 744 | ML | 9 | SC | 2 | ZM | 65 |
| CN | 57,590 | HT | 3 | MM | 28 | SD | 53 | ZW | 42 |
| CO | 945 | HU | 1,271 | MN | 31 | SE | 5,813 | | |
| CR | 57 | ID | 909 | MQ | 11 | SG | 1,877 | | |
| CU | 74 | IE | 1,791 | MR | 4 | SI | 529 | | |
